# Sub-millimeter fMRI reveals multiple topographical digit representations that form action maps in human motor cortex

**DOI:** 10.1101/457002

**Authors:** Laurentius Huber, Emily S. Finn, Daniel A. Handwerker, Marlene Bönstrup, Daniel Glen, Sriranga Kashyap, Dimo Ivanov, Natalia Petridou, Sean Marrett, Jozien Goense, Benedikt A. Poser, Peter A. Bandettini

**Affiliations:** NIMH, NIH, Bethesda, MD, USA; Maastricht Brain Imaging Centre, Maastricht University, Maastricht, The Netherlands; NINDS, NIH, Bethesda, MD, USA; University Medical Center Utrecht, Center for Image Sciences, Utrecht, The Netherlands; School of Psychology, and Institute of Neuroscience and Psychology, University of Glasgow, Glasgow, UK

**Keywords:** high-resolution fMRI, motor cortex, cortical layers, cortical columns, VASO

## Abstract

The human brain coordinates a wide variety of motor activities. On a large scale, the cortical motor system is topographically organized such that neighboring body parts are represented by neighboring brain areas. This homunculus-like somatotopic organization along the central sulcus has been observed using neuroimaging for large body parts such as the face, hands and feet. However, on a finer scale, invasive electrical stimulation studies show deviations from this somatotopic organization that suggest an organizing principle based on motor actions rather than body part moved. It has not been clear how the action-map organization principle of the motor cortex in the mesoscopic (sub-millimeter) regime integrates into a body map organization principle on a macroscopic scale (cm). Here we developed and applied advanced mesoscopic (sub-millimeter) fMRI and analysis methodology to non-invasively investigate the functional organization topography across columnar and laminar structures in humans. We find that individual fingers have multiple mirrored representations in the primary motor cortex depending on the movements they are involved in. We find that individual digits have cortical representations up to 3 mm apart from each other arranged in a column-like fashion. These representations are differentially engaged depending on whether the digits’ muscles are used for different motor actions such as flexion movements like grasping a ball or retraction movements like releasing a ball. This research provides a starting point for noninvasive investigation of mesoscale topography across layers and columns of the human cortex and bridges the gap between invasive electrophysiological investigations and large coverage non-invasive neuroimaging.

## Introduction

The human repertoire of motor activity spans an immense variety of movements, and most of our interactions with the environment involve some degree of movement. Voluntary movement is controlled by the central nervous system, particularly the brain’s motor cortex. The primary motor cortex is organized according to the principle of somatotopy, whereby areas controlling different body parts are arranged in a predictable order. Penfield and colleagues (1937) were the first to report the medial-to-lateral leg-to-face somatotopic “homunculus” in the human primary motor cortex; a representation that has been confirmed with neuroimaging for large body parts (Schieber 2002; Chainay 2004) and digits (Siero 2014; Olman 2012; Schellekens 2018) in the spatial regime of centimeters. However, fundamental deviations from this simple linear arrangement of body parts have been reported at a more microscopic level (Barinaga 1995; Strother 2012; Schieber 1993; Penfield 1937; Strick and Preston 1982; Sanes 1995; Meier and Aflalo 2008; Idovina and Sanes 2001; Hlustik 2001; Woolsay 1979; Lemon 1988). First, individual neurons in the primary motor cortex are not tuned to individual body parts but show a gradual overlap of multiple body parts (Schieber 1993, 2002; Indovina 2001). Second, the progression of body parts from face representation towards leg representation does not follow a simple steady linear order. In monkeys, multiple unconnected body part representations were reported (Kwan 1978; Park 2001). As such, it was suggested that the hand is emphasized in a core region, and the wrist, arm, and shoulder are emphasized in a half ring surrounding the core (Strohter 2012; Strick and Preston 1982; Porter and Lemon 1993; Meier and Alflalo, 2008).

In light of these deviations from a linear somatotopic organization, alternative organizational principles for the motor cortex have been investigated. Previous findings show that when a small part in the monkey motor cortex is electrically stimulated, the resulting movement combines joints and muscles in a manner resembling a coordinated action (Graziano et al. 2002), suggesting that the motor cortex organization might follow an action map representation rather than a body part map (Graziano 2016; Sanes 1995). More recently, Dietrichsen et al. also found fMRI evidence, which confirms that large portions of the motor cortex represent complex muscle synergies in humans too (Ejaz 2015). These findings are in agreement with the functioning of so called corticomotoneuronal cells that have connections to multiple muscle groups to facilitate represent complex movement actions (Omrani 2017). These corticomotoneuronal cells are, however, confined to the evolutionary younger subdivision of M1, the Brodmann area BA4p which is the inferior portion of M1 on the pre-central bank of the central sulcus (Rathelot and Strick 2006, 2009; Lemon 2008). For the evolutionary older part of M1, BA4a, it is not clear, how the action-map representation on the microscale is integrated into the body map representation on the macroscopic scale. Due to recent methodological advances in fMRI (Polimeni 2010; De Martino 2015; Huber 2017) the mesoscopic regime of sub-millimeter resolutions becomes accessible for non-invasive functional neuroimaging across columnar and laminar directions in humans. Thus, mesoscopic fMRI could provide insights on how (microscopic) action maps are integrated into (macroscopic) somatotopical body part maps. However these current mesoscopic fMR-methods can only capture small patches of cortex with thick slices and without the ability to resolve columnar structures and laminar structures simultaneously. Thus that they do not allow the investigation of microscale topographic movement representations in humans.

This study develops advanced high-resolution imaging methods and novel analysis methodology to investigate topographical organization patterns across the ‘columnar’ and ‘laminar^1^’ cortical dimensions with CBV-senitive fMRI across large patches of cortex. The method advancements could be achieved with conventional 7T MRI hardware and a non-invasive VASO (vascular space occupancy) sequence (Lu 2003; Hua 2013; Huber 2015). We investigated the neuroscientific applicability of the new methodology by mapping the meso-scale (sub-millimeter) topographical representations of the primary motor (M1) and primary sensory cortex (S1). We use this approach to investigate the organizational principle of individual finger representations for a set of simple motor actions including tapping, grasping (flexion) and retraction movements (extension). We find mesoscopic topographic finger representations in M1 (BA4a) on the order of 1.2±0.4 mm. We also find evidence of a simple topographical organizational principle: Fingers are represented in multiple somatotopically organized patches that are preferentially activated for either grasping or retraction movement actions.

## Results

To investigate the columnar organization of the primary motor cortex with fMRI, five different experiments (Fig. S1) were conducted. These were:

A. Individual tapping of two fingers (index and little).
B. Individual tapping of four fingers (index, middle, ring, and little; no thumb) interspaced with rest periods.
C. Individual tapping of all five fingers (index, middle, ring, little, and thumb) interspaced with rest periods.
D. Alternating grasping or retraction of a rubber ball interspaced with rest periods. These tasks engage muscle extension and muscle flexion of every finger.
E. Resting-state.

Motor tasks were performed with the left hand, while imaging the primary sensorimotor cortex on both sides of the central sulcus in the right hemisphere (Fig. 1A). The right hand was not engaged during any of experiments of this study. For more background on these tasks and explanations, why this task setup was used, see the supplementary information (Fig. S1). Five participants underwent 42 fMRI total sessions of 2 hours each (84h of scanning, Tab. S1). We simultaneously measured changes in the cerebral blood volume (CBV) and blood oxygenation level dependent (BOLD) response using the SS-SI-VASO method (Huber 2014; Lu 2003) with a 3D-EPI readout (Poser 2010) at 7T. The nominal resolution was 0.79 mm in the columnar dimension and 0.99 mm thick slices perpendicular to the precentral bank of the central sulcus – also known as the hand knob (Fig. 1A). Estimates of cortical depths (layers) and cortical distances (columns) across the entire central sulcus were calculated by means of simultaneously acquired anatomical and functional image contrasts in EPI space (Fig. 1B). Details on the data acquisition and analysis approach are provided in the Supplemental Experimental Procedures (Fig. S1-S10). Brain activity changes are estimated as the percent signal change of the fMRI signal during the task, compared to rest. Relative digit dominances are estimated as the activity change for a given finger compared to the activity change for all other fingers.

**Figure 1.**
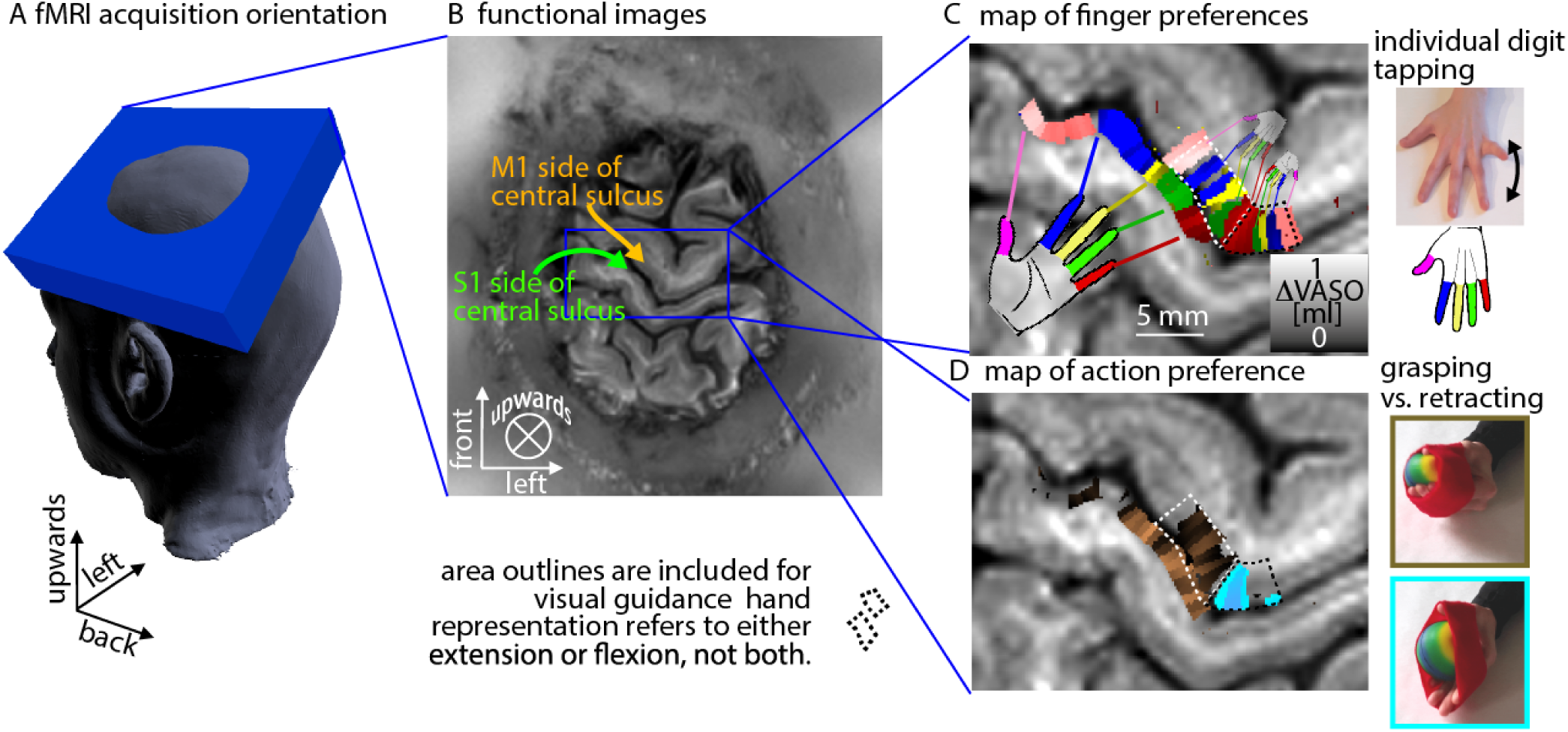
Acquisition and analysis methods for measuring the laminar and columnar functional topography. A-B) The imaging slab is aligned to cover the entire primary sensorimotor cortex and slices are tilted as indicated by the blue box through the right cortical hemisphere. C) The functional CBV signal changes obtained using a five-finger tapping task reveal individual digit representations in M1 and S1. While S1 has a homunculus like linear representation of individual digits, the primary motor cortex shows clear deviations of the homunculus model. D) While S1 has a stronger activity for grasping compared to retraction motor actions, the primary motor cortex shows individual patches of areas that are either specific to grasping or retraction actions. For further graphical guidance about the respective dimensions and coordinate systems, see Fig. S3. All functional results shown here refer to left hand movement tasks. The right hand was not engaged here.

Functional MRI signal changes in S1 show a linear arrangement of individual digits (Fig. 1) as previously described in high-resolution fMRI (Olman 2012; Panchuelo 2016; Kolasinsky 2016; Schluppeck 2017; Siero 2014; Ejaz 2015). The distance between the representations of the thumb and the little finger in S1 is 16 ± 4 mm. In M1 we find clear deviations from a continuous linear alignment of the digit representations; we find multiple representations of every digit in a mirrored pattern (Fig. 2). These digit representations are significantly smaller than the representations in the sensory cortex. The distance between the representations of the thumb and the little finger in M1 is 6 ± 2 mm. The mirrored pattern of multiple distant digit representations was highly consistent across participants (Fig. 2), across fMRI contrasts (Fig. S5), and across days (Fig. S6).

**Figure 2.**
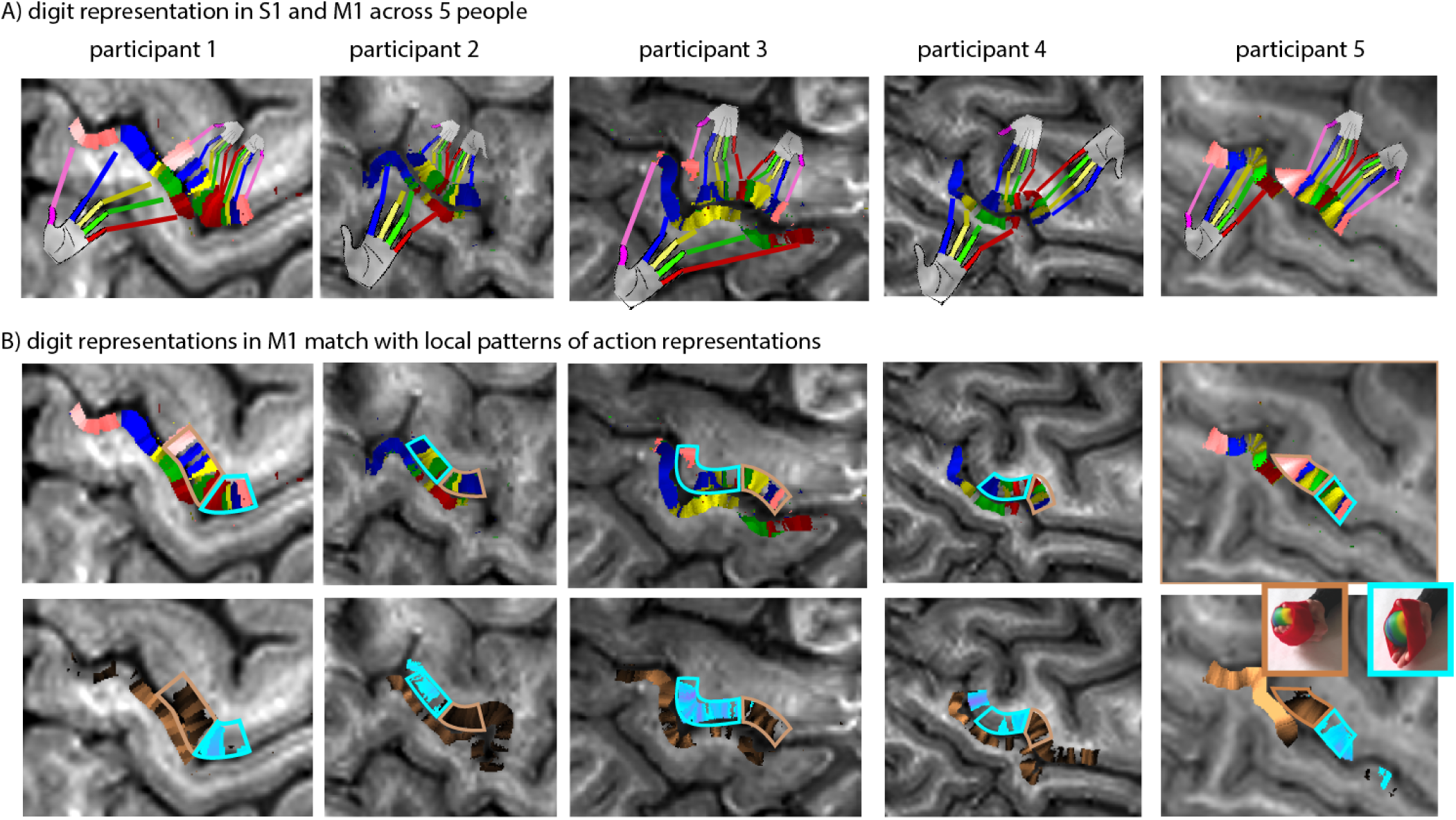
Multiple hand representations across participants. A) The finger dominance maps show across all participants that the primary sensory cortex has a single representation of each finger in all participants, while the primary motor cortex has multiple finger representations of each finger. The representations in the primary motor cortex are mirrored and about half the size of the representations in the primary somatosensory cortex. B) The spatial pattern of multiple finger representations is compared to the representations of different movements, grasping and retraction of a ball. Each complete set of fingers is outlined with manually drawn borders. These borders match the outlines of the different action tasks. The copper and turquoise colors refer to preferred grasping and retraction preference, respectively. The functional contrast refers to relative action preference between task conditions. I.e., turquoise patches refer to stronger activity during retraction periods compared to grasping periods (see also Fig. S7 for absolute signal changes). The maps in the left column are the same as in Fig. 1. All functional results shown here refer to CBV changes during left hand movement tasks. The right hand was not engaged.

We find that different patches of the hand knob in M1 are preferentially activated during grasping and retraction movements, respectively. These patches are 6 ± 2 mm in size and columnar distance. We find that the outlines of grasping and retraction preference patches contain complete sets of all digit preferences (copper and turquoise outlines in Fig. 2B).

We next quantified relative preferences for body parts (i.e., different fingers) and movements (i.e., grasping versus retraction) along the medial-lateral axis in both the primary motor and primary sensory cortex. Columnar profiles along the central sulcus are shown in Fig. 3, for one representative participant. In the primary motor cortex, we find a sharp switch in preference between grasping and retraction at a certain point along this axis (Fig. 3A left, echoing results shown in the last row of Fig. 2). In the primary sensory cortex, we find a strong preference to grasping tasks only. There are no clear areas that have a stronger S1 response to retraction versus grasping (Fig. 3A, right). This is likely because common grasping actions are more often associated with stimulations of the sensory receptors (exteroception on the fingertips and inside of the hand) compared to retraction actions. Representations of sensory proprioception in the primary sensory cortex might be similarly engaged for grasping and retraction movements and, thus, do not introduce a preference to either of the two motor actions in the sensory cortex. The columnar profiles in Fig. 3B also clearly demonstrate the multiple mirrored finger representations in M1, versus the single linear representation in S1. Finally, we quantified relative finger preferences across both columnar and layer dimensions (Fig. 3C). We find that there is a more gradual transition between preferred fingers (i.e., more overlap across finger representations) in superficial layers (solid lines) than in deeper layers (dashed lines), where the transition is sharper. This result is also confirmed in resting-state data (Fig. S8).

**Figure 3.**
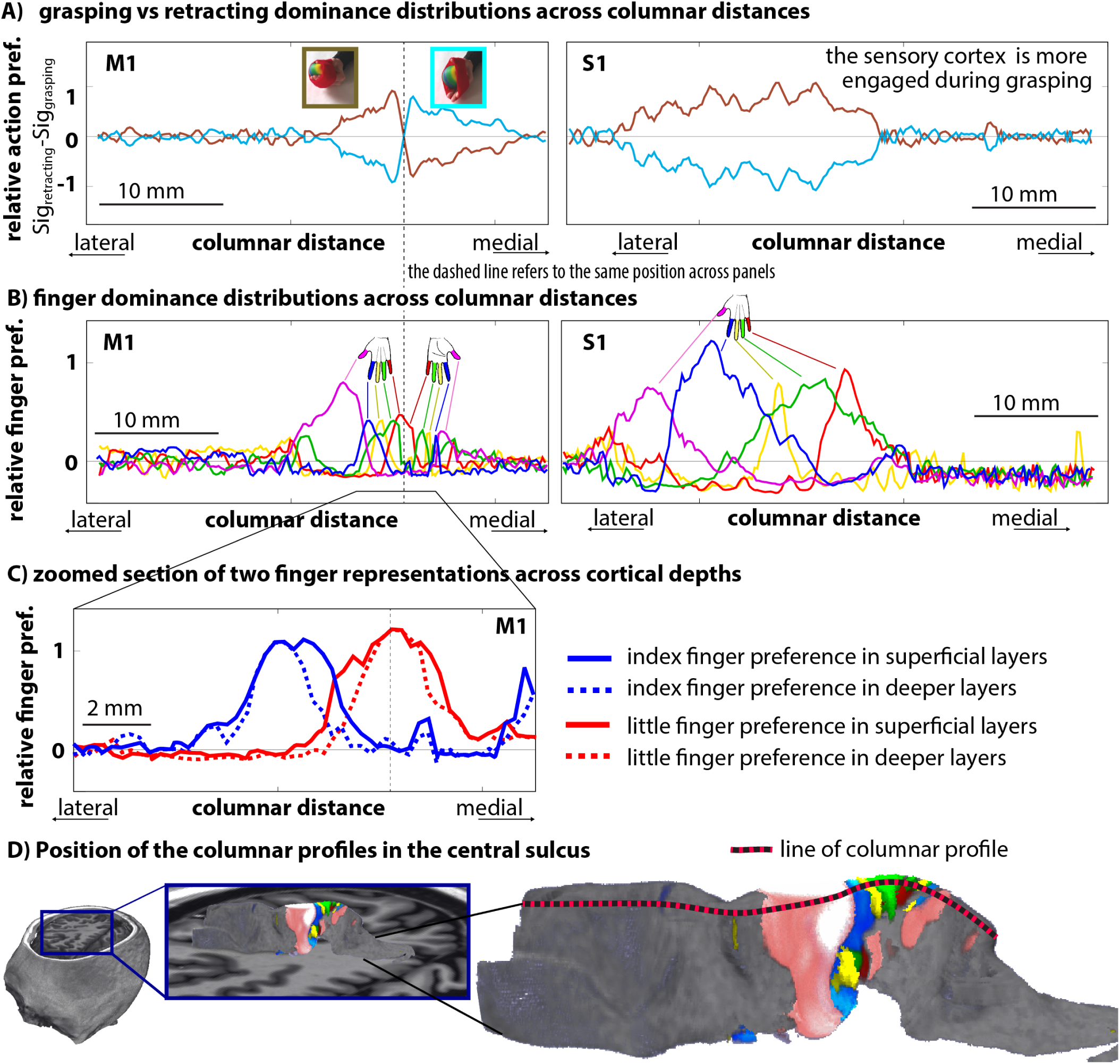
Columnar profiles of functional representations along the central sulcus of participant 1. Panel A) depicts the columnar distribution of grasping preference and retraction preference. While the entire primary sensory cortex is more engaged for grasping actions, the primary motor cortex has distinct patches that are preferentially more active for either grasping or retraction. Panel B) shows that each of the patches in the primary motor cortex contains a complete set of linearly aligned finger patches. Panel C) shows that superficial cortical layers have larger overlap across fingers compared to deeper layers. The dashed line in panel A-C refers to the same columnar position. The unit of the y-axis refers to the relative signal change between conditions (see Fig. S5). The results refer to a projection of a 0.79 mm slab through the central sulcus (D). The scale bars refer to the spatial distance along the curved GM accounting for the average curvature in the projection slab. The anatomical contrast along these columnar profiles is given in Fig. S7 for reference.

Besides task-based topographical descriptions, we also characterized the topography of finger representations in restingstate functional connectivity with the sensory cortex. The aim of these investigations was to confirm the multiple digit representations in the motor cortex on data that does not rely on a specific task design. As expected from the task-based data, we find the same multi-stripe patterns in the primary motor cortex for A) task-induced motor activity, B) resting-state seed-based correlation, and C) selected independent components of FSL-melodic ICA (Fig. 4). In all three cases, the two investigated fingers (index and little) are represented as two clear patches in the sensory cortex (red and blue arrows in bottom left of Fig. 4A-C). In the motor cortex, however, multiple red and blue patches can be identified. The resulting stripe-pattern of blue-red-blue is very similar for task and resting-state results (red and blue arrows in top right of Fig. 4A-C). We find that the digit representations in the sensory cortex have a stronger connectivity with the grasping patches in the motor cortex compared to its retraction patches (Fig. 4E-D). This is consistent with the task results showing that the sensory cortex has stronger activity for grasping motor actions compared to retraction motor actions.

**Figure 4.**
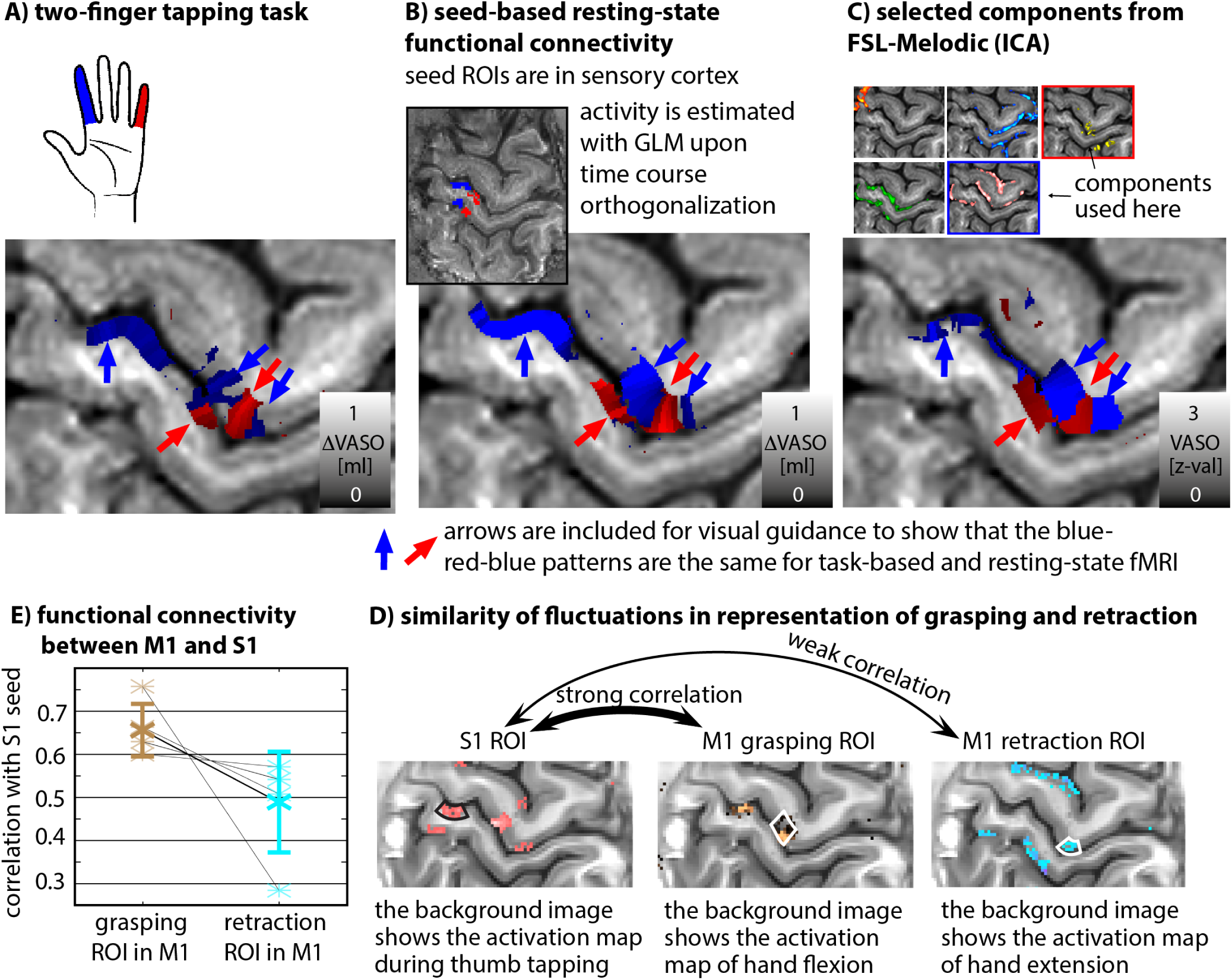
Patterns of similar temporal fluctuations in resting-state activity match the patterns of task-evoked activity. A) Map of voxelwise preferences to become active during index finger movement or little finger movement. As shown in Fig. 1-2, the primary motor cortex has multiple representations of individual fingers. B) Map of resting-state signal fluctuation similarity for seed regions in the primary sensory cortex (inset). The colors refer to seed regions of interest (ROIs) that represent the little finger and index finger in S1. C) Very similar networks are generated using selected components from an ICA decomposition as seeds for connectivity analysis. Here we selected two (the ones outlined in blue and red) and used those as “seeds” to compute a preferential correlation map. D) The functional resting-state connectivity between individual finger representation areas in M1 and S1 are stronger for regions that represent grasping actions than for regions that represent retracting actions. E) Seed-based correlation analysis between the thumb representation in S1 and two thumb representations in M1 reveal that the flexion representation has larger functional connectivity to S1 than the extension representation. This might be associated with the fact that common grasping actions are more often associated with stimulations of the sensory receptors (on the inside of the hand) than retraction actions. The star symbols refer to individual participant and the error bars refer to the inter-participant standard deviation.

## Discussion

The data presented here suggest a mesoscopic topographical organization principle of individual fingers in the primary motor cortex (BA4a) at an unprecedented spatial resolution. The primary motor cortex has been extensively studied with cytoarchitecture, functional representations using invasive electrophysiology, human non-invasive electrophysiology and neuroimaging. No comprehensive view, however, has emerged regarding how the representations of actions, as found with electrical microstimulation (Strick and Preston 1928, Graziano 2002, 2016) translate into topographical somatotopic organization principles, as detected for larger body parts. Here, using high-resolution CBV-fMRI neuroimaging in humans and topographical analysis tools, we reveal how these different organizational principles are integrated across cortical columns and layers.

We find evidence for action specific representations of fingers in M1, as well as for finger specific representations. Our results reveal how these multiple organizational principles are elegantly combined in M1: We find multiple mirrored representations of individual fingers that are differently engaged for specific movement actions (Fig. 1, 2, 3). Thus, M1 is organized as an action map as well as a topographical finger map in small (0.9-1.2 mm) representations along the columnar dimension within 4-9 mm action patches.

The multiple representations of individual fingers can be seen in both task-based fMRI and resting-state functional connectivity with S1 (Fig. 3). We find that the finger representations in the motor cortex are very similar across cortical depths. There are only small differences between superficial cortico-cortical input layers II/III and corticospinal output layers Vb/VI. Namely, the size of the finger representations is slightly smaller in deeper layers compared to superficial layers. One potential origin of the sharper body part representation in deeper layers might be associated with previously described phenomena of surround inhibition in M1. This kind of surround inhibition in M1 has been proposed to serve the function of improved discriminiability between motor representations. The phenomenon has been described as the reduction of corticospinal excitability in muscles that are non-active but adjacent to the active muscles (Beck and Hallett, 2011). During voluntary finger flexion, the person’s muscle shows increased corticospinal excitability, whereas the surrounding neighboring muscle shows decreased corticospinal excitability. However, due to the lack of invasive layer-dependent electrophysiology measurements in the literature about M1, it is unclear whether this is expected to be different across cortical layers, as it is in sensory brain areas (Hubel and Wiesel, 1968, 1972). In resting-state fMRI, we find that the functional connectivity strength varies, as it also does with task-based fMRI, with more tightly constrained connectivity in deeper layers compared to superficial layers (see Fig. S8).

As opposed to previous columnar fMRI studies, we do not only try to depict known structures with known shape and size as proof-of-principle for a method as previous studies. Instead here, we are finding previously unknown organization principles of sub-millimeter representations in M1. This is a fundamentally new approach and a paradigm shift for the field of *columnar* and *laminar* fMRI.

It is important to note that the motor cortex representations of grasping vs. retraction are not completely orthogonal to each other: that is, the entire hand knob shows increased activity to both grasping as well as extension movements, compared to rest. The respective grasping and retraction patches show differential sensitivity to either motor action (Fig. S6C) only. Other potential movements could theoretically engage the two patches even more deferentially: for example, spreading fingers vs. clenching fist; maintaining force without movement (i.e., isometric muscle contraction) vs. movement with minimal force, and others. These potential movement actions are only partly segregated for the particular flexion vs. extension task used here. For the finger tapping tasks used here, the individual finger movements underwent both extension and flexion movements. Thus, the double-hand map can relate to grasping and retraction task that engages flexion and extension differently.

In the primary somatosensory cortex, we find no deviations from the homunculus model as shown previously in humans (Schluppeck 2017; Olman 2012; Kolasinski 2016; Shellekens 2018).

Previous digit mapping studies using GE-BOLD fMRI could not clearly identify the mirrored finger representation in the primary motor cortex. This might be due to the fact that GE-BOLD fMRI suffers from poor localization specificity due to the presence of large draining veins (Turner 2002; Menon 2002; Kim and Ogawa, 2012; Kennerley 2015). Most human fMRI studies that investigated somatotopic organization likely did not have the sub-millimeter specificity to investigate deviations from the linear representation without methodological challenges (Olman 2012; Siero 2014; Dechent and Frahm 2003, Hlustik 2001; Shellekens 2018). We believe the columnar specificity of GE-BOLD could be limited by two potential sources of venous contaminations. First, large principal veins are described to have tangential branch lengths of 2-6 mm (Fig. S8E,; Duvernoy 1981). Thus, these tangential veins can result in more tangential signal mixing of GE-BOLD signal compared to their respective neural representations. Second, in some participants the sensory and motor banks of the central sulcus can be drained by the same pial veins. BOLD signal changes in the precentral gyrus can thus be contaminated by signal leakage from the postcentral gyrus. Other methodological challenges to identify the double-hand signature may arise from sensitivity limitations, task design, analysis design and effective resolution. Our results show that CBV-based fMRI has a higher localization specificity than conventional GE-BOLD fMRI (Fig. S2, S5), thus overcoming the specificity limitations of previous studies. Even though CBV-based fMRI has a lower sensitivity and requires longer scan durations to exceed the detection threshold, it provides clearer results. The higher localization specificity of CBV-fMRI has previously been shown in humans with VASO fMRI (Huber 2017) across cortical layers. In this study we extend this finding and also show the higher localization specificity of VASO across cortical columns. Note that the CBV weighting in VASO has been extensively validated by comparisons with gold-standard methods in rats and monkeys across layer and columns (Huber et al., 2015a-c; Kennerley et al., 2013).

To optimally separate individual digit representations, we included long rest periods into the experiment and used a pseudo random ordering of individual fingers. This is different to previously employed “phase-encoding” paradigms and it comes at the cost of less efficient task design requiring longer scan durations. However, it can be more sensitive to detect deviations of continuously linearly aligned representations.

Most of the knowledge on the functional representation of movements in the primary motor cortex has been obtained from countless experiments in monkeys over the last century. The current state of consensus in the field is nicely summarized by Paul Cheney in (Omrani 2017; see also references therein); Overall, corticomotoneuronal cells in the primary motor encode muscle-related parameters of movement such as muscle activity and muscle force. Although some corticomotoneuronal cells in the primary motor cortex (particularly those involved with finger movements) have their terminations confined to motoneurons of single muscles, a large amount of corticomotoneuronal cells are not rigidly coupled to the activity of its target muscles but show specialization for particular movements or categories of muscle activity. Namely, almost half of the corticomotoneuronal cells facilitate muscles involving at least one distal and one proximal joint and are specialized for specific muscle synergies, E.g. for reach-to-grasp movements. With respect to action representations shown in Fig. 2B, it is important to note that Cheney and Fetz (1985) had previously identified the muscle fields of neighboring corticomotoneuronal cells. They showed that neighboring corticomotoneuronal cells had muscle fields that were very similar. Hence, the notion of cortical patches that are preferentially activated for grasping and retraction actions, as shown in Fig. 2B, has its basis in previous monkey data and could refer to these previously described muscle fields.

A previous study by Ejaz et al. (2015) already identified deviations from linear organizations for finger representations in the human motor cortex with GE-BOLD at 2.5 mm and 1.4 mm resolutions, respectively. These data already showed some indications of multiple finger representations (e.g. Fig. 1 in (Ejaz et al. 2015)). However, these data were not discussed with respect to an alternative geometric somatotopic organization principle such as a mirrored representation.

Previous studies with PET had shown multiple hand movement representations across both sub-areas of the primary motor cortex (Geyer 1996), BA4a and BA4p, respectively. The mirrored double-hand representation shown here (Fig. 2-3) is located in the anterior side (BA4a) of the primary motor cortex, only. Thus, the hand representations in BA4p, as discussed in (Geyer 1996) refers to yet another representation of the fingers. This is the evolutionary younger part of M1 that is located deep in the central sulcus. In this part of M1, individual body parts are largely overlapping (probably to facilitate complex hand movement) and thus, in this part of the motor cortex finger dominance maps might be misleading ways of depicting the complex representation principle (Ejaz et al., 2015). We find that this finger representation in BA4p is actually not completely separated from the one in BA4a. It is partly connected to the representation in BA4a and does not show a mirrored pattern (Fig. S9).

In contrast to human fMRI literature, electrophysiological recordings have shown deviations from a linear somatotopic organization. However, much of our understanding from the electrophysiological organization in M1 comes from experiments in which stimulations or recordings are performed from few cortical points only, whereas neuroimaging samples continuously and uniformly across space. Thus, continuously sampled maps of the motor cortex representations with electrophysiology at the sub-millimeter resolution have not been collected across columns and layers yet. And consequently, most of these studies only conclude that the representation pattern is complicated and nonlinearly organized (Schieber 1993; Hatsopoulos 2010) without proposing an alternative organization principle.

In this study we show for the first time that individual fingers have multiple representations in a mirrored pattern along the lateral-medial axis of primary motor cortex, with each whole-hand instance corresponding to a distinct movements. This mirrored representation across grasping and retraction patches gives rise to neighboring representations for movement synergies (d’Avella and Bizzi 2005). As already suggested by Penfield (1937), our data show that the Penfield homunculus model is an oversimplification. With high resolution fMRI, we can confirm deviations of the homunculus model in M1 at very high resolutions corresponding to small body parts. Our data agree with the hypothesis that in the submillimeter regime, the motor cortex is organized as an action map (Graziano 2016).

The present data corroborate several findings from invasive electrophysiologic recordings and microstimulation experiments in rodents and monkeys, that have elucidated the organizational principle of M1, and extend those to the level of the human M1. The presented methodology allows noninvasive recording of mesoscale functional representations in human M1 that were previously inaccessible. This might help to further understand the blueprint of cortical control over movement dynamics and kinematics. Beyond that, functional imaging of M1 at the layer and columnar level may help to elucidate how aberrations in the organizational principle could lead to movement disorders with largely unknown pathophysiology, such as dystonia.

## Summary

Here we used advanced non-invasive neuroimaging in humans to provide novel insights in the organisational principles of the primary motor and sensory cortices at the mesoscale. We demonstrate for the first time that individual fingers are represented multiple times in the primary motor cortex in a columnar fashion following a mirrored pattern, and these representations are differentially engaged during specific motor actions (i.e., grasping versus retraction movements). By using new imaging and analysis technology that bridges the gap between invasive electrophysiological recordings and non-invasive large coverage mesoscopic fMRI in humans, we resolve previous controversies of M1 representation principles as ‘body map’ at the macro-scale vs. ‘action maps’ at the micro-scale.

## Acknowledgements

The research was supported by the NIMH Intramural Research Program (ZIA-MH002783). We thank Kenny Chung and Harry Hall for radiographic assistance. The study was approved under NIH Combined Neuroscience Institutional Review Board protocol 93-M-0170 (ClinicalTrials.gov identifier: NCT00001360). Laurentius Huber was funded form the NWO VENI project 016.Veni.198.032 for part of the study. Portions of this study used the high-performance computational capabilities of the Biowulf Linux cluster at the National Institutes of Health, Bethesda, MD (biowulf.nih.gov). We thank Hartwig Siebner and Mark Hallett for comments on the manuscript. We thank James Kolasinsky for discussions about intermediate results of this study and task design. This preprint is formatted based on a LATEXclass by Ricardo Henriques.

## Author contributions

L.H., conducted the experiments. L.H., P.B. designed the tasks. L.H., E.F. designed the analysis. L.H., S.K., D.G. wrote the analysis code. B.P., L.H., D.I. designed the MR-sequence, D.H., M.B., N.P., S.M., J.G. provided advice on experimental design, sequence, analysis, and research direction. L.H. wrote the draft of the manuscript. All authors contributed to the study story line and edited the manuscript.

## Data and Software availability

All raw, anonymized MRI data of this study can be anonymously downloaded (https://goo.gl/1rdfLe). Fully processed data of two participants can be downloaded (https://activecho.cit.nih.gov/t/9ewfvz13). All custom written software (source code) and evaluation scripts are available on Github (https://github.com/layerfMRI/Tolopoly_strips). The authors are happy to share the 3D-VASO MR sequence upon request via a SIEMENS C2P agreement. A complete list of scan parameters used in this study is available on Github (https://github.com/layerfMRI/Sequence_Github/tree/master/Topology).

## Supplementary methods

### Sample size

Five healthy right-handed volunteers (age 23, 25, 30, 33, 34 years) participated in this study. The number of participants was chosen based on previous comparable studies (Huber 2015a; Muckli 2015)^2^. Every participant underwent at least six two-hour MRI sessions. A list of all performed scans that are available to download is given in Tab S1.

**Table S1.** Number of scans acquired in this study that are available to download. Five volunteers participated in this study. Every participant underwent 5-9 functional scan sessions and 3-10 additional anatomical sessions.

|  | subj 1 | subj 2 | subj 3 | subj 4 | subj 5 |
| --- | --- | --- | --- | --- | --- |
| two | 1 | 1 | 1 | 1 | 1 |
| four | 2 | 1 | 1 | 1 | 1 |
| five | 2 | 2 | 0 | 1 | 1 |
| ext vs. flex | 2 | 1 | 1 | 1 | 1 |
| rest | 3 | 2 | 1 | 1 | 1 |
| 0.5 anat | 3 | 3 | 3 | 3 | 3 |
| 0.7 anat | 10 | 10 | 7 | 6 | 3 |

Five additional participants were invited for pilot experiments to optimize parameters for the MRI-sequences, task design and reconstruction parameters (See Fig. S10). An additional 3 participants were invited for pilot experiments to explore the best set of GRAPPA parameters (Kernel size, algorithm type, regularization strength) for image reconstruction.

### Imaging hardware

In this study, we used standard imaging hardware commonly available in more than 50 high-field imaging centers, a MAGNETOM 7T scanner (Siemens Healthineers, Erlangen, Germany) using the vendor-provided IDEA environment (VB17A-UHF). The scanner was equipped with a SC72 body gradient coil (maximum effective gradient strength used here: 47 mT\m; maximum slew rate used: 190 T\m\s). For RF transmission and reception, a volume-transmit\32-channel receive head coil (Nova Medical, Wilmington, MA, USA) was used.

### Tasks

Five different functional tasks were employed across multiple fMRI scan sessions in this study. Schematic depictions of the tasks are given in Fig. S1.

We used multiple, partially redundant experiments that involved the same digits. Our reasons for some choices of the task design were:

- Having multiple tasks with a varying number of fingers involved, provided us with a good spectrum of sensitivity and specificity. As such, experiment A) had 38 averages for each task condition and thus offers highly sensitive results. But since only two digits were involved, the spatial specificity was limited. Experiment C, on the other hand, included only 8 averages of each task condition and thus yields less sensitive results. However, since all five fingers were engaged, the spatial specificity was higher.
- It was not clear that the co-registration across different experiments on different days with different shims can be achieved the same sub-millimeter accuracy as the co-registration within one MRI session. Thus, using the same fingers across different days allowed us to investigate the consistency of the digit representation location despite technical analysis challenges and potential co-registration inaccuracies.
- For some of the resting-state analyses, it was highly beneficial to restrict the analysis to two seed regions only. With two seed regions, it is straightforward to orthogonalize the seed time courses to each other. This allows us to minimize the effect of shared sources of variance such as physiological noise. Therefore, we acquired a task-based reference dataset that involved only two fingers.
- Dependent on how the participant is moving the thumb during the tapping, there can be a residual risk that this movement action also involves wrist motor activity (personal discussion with Dr. James Kolasinski), which is not the case for the index finger. To avoid the risk of wrist activity during thumb tapping, we refrained from using thumb and little finger for the two finger tapping task. Instead, we restricted the two-finger tapping task to index/little finger. Even though they are expected to be represented spatially closer to each other in M1, they are better comparable with respect to the degree of individualization and typical hand movement synergies (Thakur 2008).
- In task B) and C), the rest periods are not equally distributed between every individual finger. This was done in order to allow more trials per finger in a given experimental duration of 34 min, at the cost of potential signal leakage across digit representations. The restperiods are pseudo randomly omitted between consecutive finger tappings. Here, omitted rest periods are chosen in a way that the response of pairs of fingers can be resampled to match the results of other experiments with fewer engaged fingers. As such, combining the responses of index/middle finger and ring/little finger in the four finger experiment should give rise to similar topographical striping pattern as the two finger experiment. This was helpful as a consistency check across days and tasks.

### MRI acquisition

The timing of magnetization preparation and interleaved acquisition of VASO and BOLD data was done as previously described (Huber 2017). In short, the timing of the acquisitions is: TI1/TI2/TR=1200/3600/4400 ms across all experiments. The adiabatic VASO inversion pulse is based on the TR-FOCI pulse (Hurley 2010), with a duration of 10 ms. Inflow of fresh blood was minimized by the application of a phase skip during the frequency sweep (Huber 2014). Online reconstruction was performed using a combination of standard scanner software with kernel size 4×3×3. Slab-oversampling was applied in the second phase direction with 9.1%. 3D slice aliasing was minimized using a sharp slab-excitation pulse profile with a bandwidth-time-product of 25. A variable flip angle distribution was used to minimize T1-blurring along the second phase encoding direction. The acquisition of the GRAPPA calibration data followed the FLASH approach as implemented in (Ivanov 2015). In order to minimize T2*-blurring, no-partial Fourier imaging was applied. This means that we were forced to use a relatively long echo time of 32 ms, trading off SNR for a higher localization specificity. The imaging matrix was 162 × 162 × 24, with a nominal resolution of 0.79 × 0.79 × 0.99 mm3. Magnitude images from individual RF receive channels were combined with the sum-of-squares method. Further sequence parameters are given in the protocol pdf of the vendor (https://github.com/layerfMRI/Sequence_Github/blob/master/Topology/Topology_VASO.pdf). As a means of finding anatomical landmarks of high myelination across columns and layers, we acquired 3-10 averages (parts of these data were acquired in combination with previous studies) of 0.7 mm whole brain MP2RAGE (Marques 2010) and 3 averages of 0.5 mm slab MP2RAGE and Multi-echo FLASH (TEs = 8.5/23.2 ms) data.

**Figure S1.**
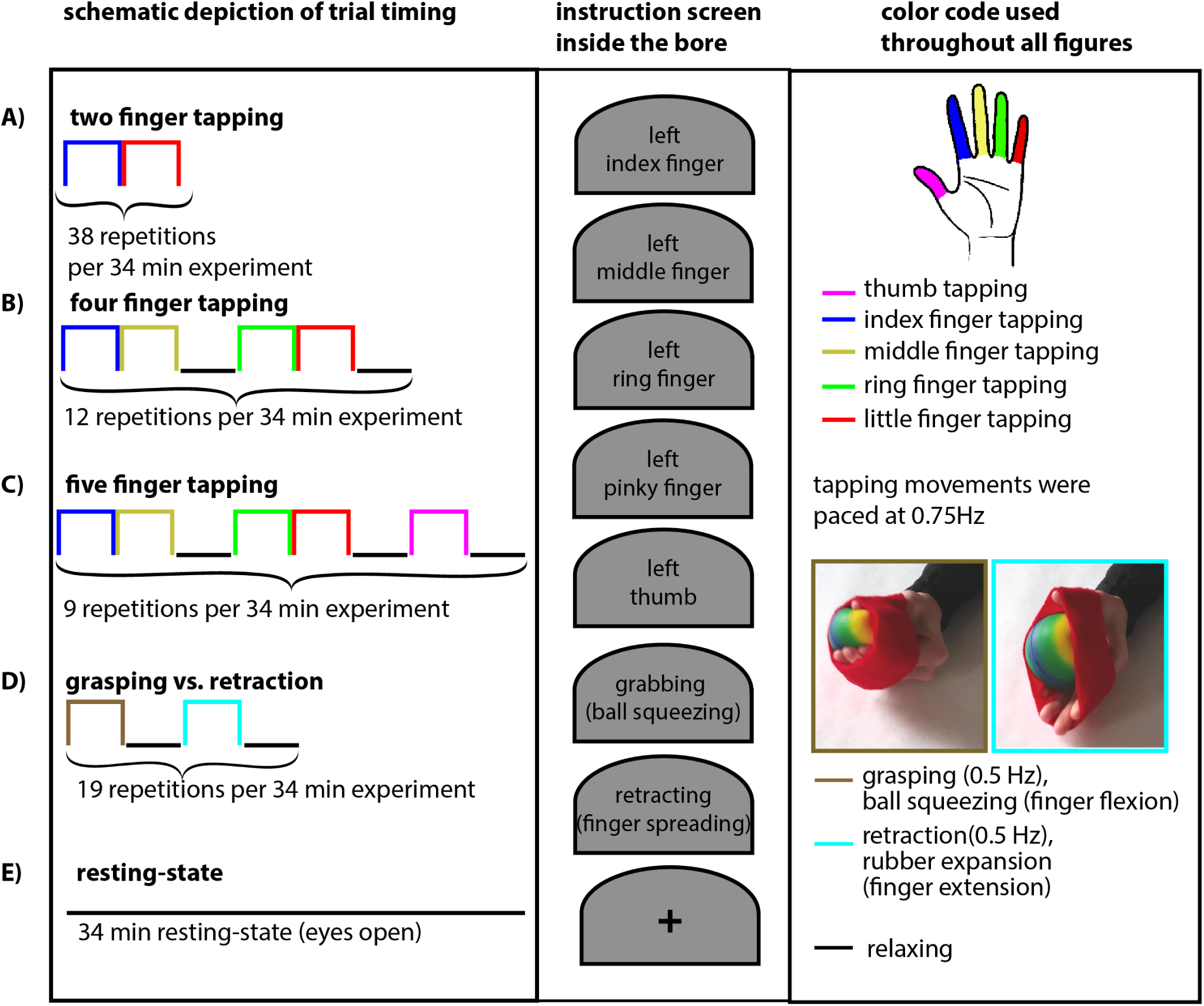
Functional tasks used in this study. A) Finger tapping of two fingers; index finger and little finger without rest periods. B) Finger tapping of four fingers; index finger, middle finger, ring finger, and little finger with randomly interspaced rest periods. C) Finger tapping of five fingers; thumb, index finger, middle finger, ring finger, and little finger with randomly interspaced rest periods. D) Grasping of rubber ball and retracting against rubber band with regularly interspaced rest periods. Panel E) represents resting-state scans. Each activity period (colorful blocks) and each inter-stimulus interval (black line) lasted 26.5 seconds. Every experiment lasted 34 min and was repeated within sessions and across days. Every task condition (tapping of one individual finger, grasping, retraction, and inter-stimulus rest) lasts 26.5 sec. Tapping movements were paced at a frequency of 0.75 Hz. Instructions of the task were given to the participants via a projector and mirror system inside the scanner bore. All movement tasks were done with the left hand. The right hand was not engaged for any task in this study.

### Analysis of functional data

All MR images were motion corrected using SPM12 (Functional Imaging Laboratory, University College London, UK). Volume realignment was done with a 4th-order spline inter-polation and realigned volumes were written out using a 4th-order B-spline interpolation. The outermost slices (one at each side) were excluded from the analysis to minimize any residual 3D-EPI related slab fold-over artifacts and reslicing artifacts. Activation GLM analysis was done using FSL FEAT (Version 5.98). Odd and even time points were treated separately and divided by each other to correct for BOLD contaminations in VASO (Huber 2014). The functional percent signal changes were calculated as 100 × (*S_activity_ – S_rest_*)*/S_rest_* (Fig. S2 left columns of panels respectively). For relative finger dominance analysis, we estimated the finger selectivity as:

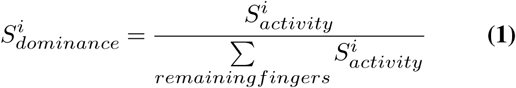

with *i ∈* {*thumb, index, middle, ring, little*}, while 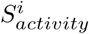 refers to the above percent signal change (middle column in panels of Fig. S2). The grasping and retraction reference was calculated as 100 *×* (*S_grasping_ − S_retraction_*)*/S_rest_* and 100 *×* (*S_retraction_ − S_grasping_*)*/S_rest_*, respectively.

Alignment across runs and sessions was conducted as part of the motion correction. To avoid the risk of spatial blurring due to additional resampling and imperfect alignment, no distortion correction was applied. Instead, all columnar and laminar analyses were performed directly in the EPI space, which is possible due to the inherent T1 contrast that arises from the inversion-recovery property of the functional VASO sequence (Huber 2017).

### Analysis across laminar and columnar structures

Boundary lines of the gray matter (GM) ribbon to cerebrospinal fluid (CSF) and white matter (WM) were estimated with FreeSurfer (v6.0) with whole-brain MP2RAGE data that had been registered to the functional EPI space with ANTS. These boundaries were used as visual guidance to manually segment GM in EPI space slice-by-slice, as described in the manual (https://layerfmri.com/getting-layers-in-epi-space/). A coordinate system across cortical layers and columns was estimated in LAYNII (https://github.com/layerfMRI/LAYNII). LAYNII is an open source C++ software suite for computing layer and column functions. We estimated the depth of equidistantly distributed layers. Note that in the context of fMRI, these estimates of cortical depth do not refer to cytoarchitectonically defined cortical layers. The cortical thickness of the primary motor cortex is 4 mm (Fischl and Dale 2000). With our resolution of 0.79 mm, we obtained 5-7 independent data points across the thickness of the cortex. Across these data points, we created 20 layers across the thickness of the cortex on a 4-fold finer grid than the effective resolution. The number of twenty layers was chosen based on previous experience in finding a compromise between data size and smoothness (see Fig. S6 in (Huber 2018) and https://layerfmri.com/how-many-layers-should-i-reconstruct/). Columnar profiles in Fig. 3 and Fig. S4 are generated from unsmoothed data. For Figs. S3 and S6, the functional signal was directionally smoothed with 0.5 mm within columns and extracted in sheets to produce a reconstruction of the face of the anterior bank of the central sulcus. No smoothing was applied in the direction across columns.

### Cortex unfolding using IMAGIRO

For efficient depicting of MRI contrast across the entire central sulcus, we used the program LN_IMAGIRO of LAYNII to unfold both GM banks of the central sulcus. Fig. S3 depicts this spatial operation schematically.

### Location of double-hand representation is compared to anatomical landmarks

It has been suggested that there are distinct septa of low myelination that provide anatomic landmarks, delineating the borders of major body parts like trunk, face and hand representations (Kuehn 2017). Such anatomical landmarks could be valuable as a reference for comparison with ex-vivo histology data (Ding 2016; Wagstyl 2018) providing insights to the interaction of structure and function. Furthermore, such anatomical landmarks could have the potential to be used as a predictor for the location of individual finger representations without functional localizer experiments. Thus, estimates of T1, T2*, and phase were extracted in the same columnar coordinate system as the functional data. Fig. S4 depicts all the anatomical contrasts that were used to identify anatomical septa landmarks. Arrows refer to respective locations in the column profiles of Fig. S4 and Fig. 3.

Based on the data shown in Fig. S4, we do not find very clear deviations of anatomical contrasts in and around the hand area that are significantly different from the overall variance across columns. However, we find some indications of changes in T1 that are overlapping with the position of the thumb and index finger representations, both in S1 and in M1 (colorful arrows in Fig. S4), which could be interpreted as indications for septa.

### Reproducibility of results across different fMRI contrasts and task paradigms

The robustness of the double-hand representation in M1 was assessed across multiple fMRI contrasts (BOLD and VASO), tasks (two-finger tapping, four and five finger tapping), as well as in resting-state functional connectivity with S1 (Fig. S5). Across all conditions, we find multiple linearly aligned finger representations in the motor cortex. It is most clearly visible in the VASO results with two fingers tapping as a blue-red-blue pattern. GE-BOLD results show similar patterns but are less clear. In the BOLD results there is more variability in finger representation size and ordering. We attribute this is to confounding venous signal leakage artifacts.

**Figure S2.**
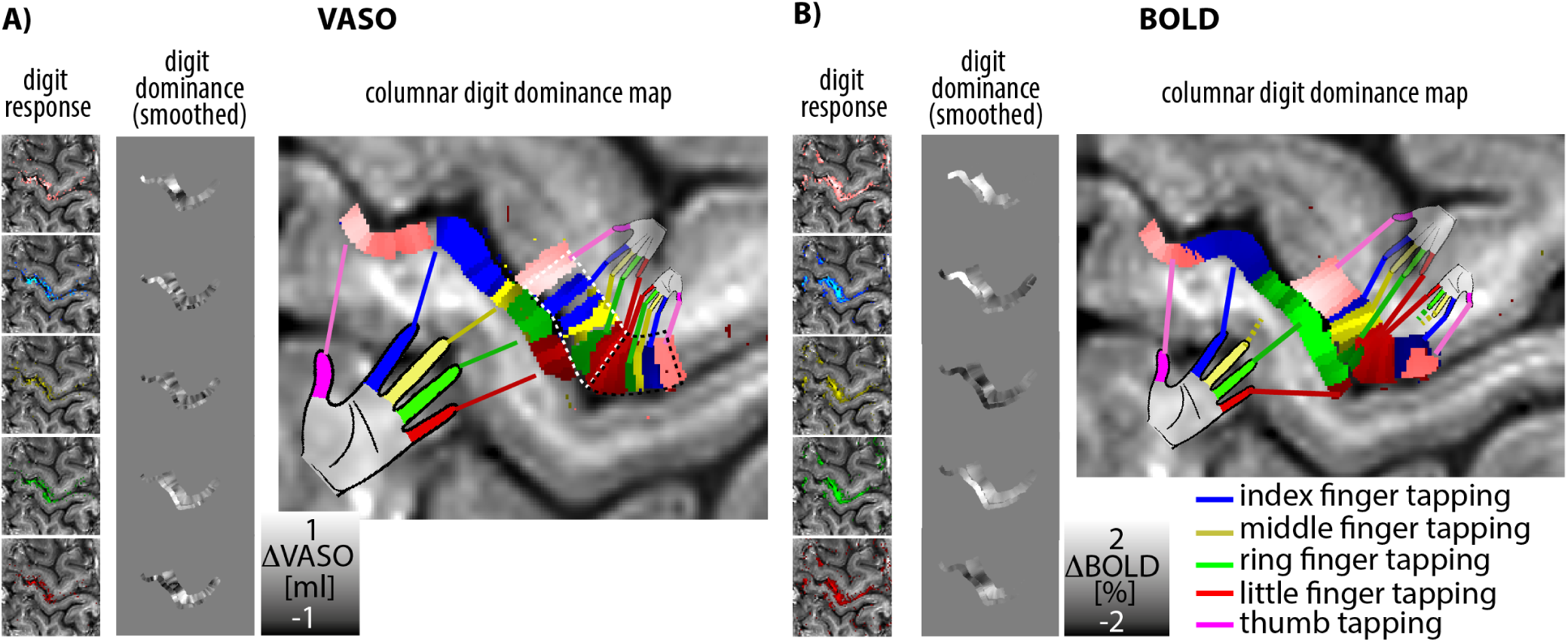
Depiction of the steps to generate digit-dominance maps. First, the percent fMRI signal change is calculated for every individual digit (left columns in panel A and B). In the next step, the finger dominance is estimated by dividing the individual digit’s response by the sum of all other digits’ responses. The resulting relative digit dominance values are directionally smoothed within columns (middle columns in panel A and B). The final topographical maps are generated by overlaying responses of all fingers with different color coding. These are the same type of maps as shown in Fig. 1C and 2A.

**Figure S3.**
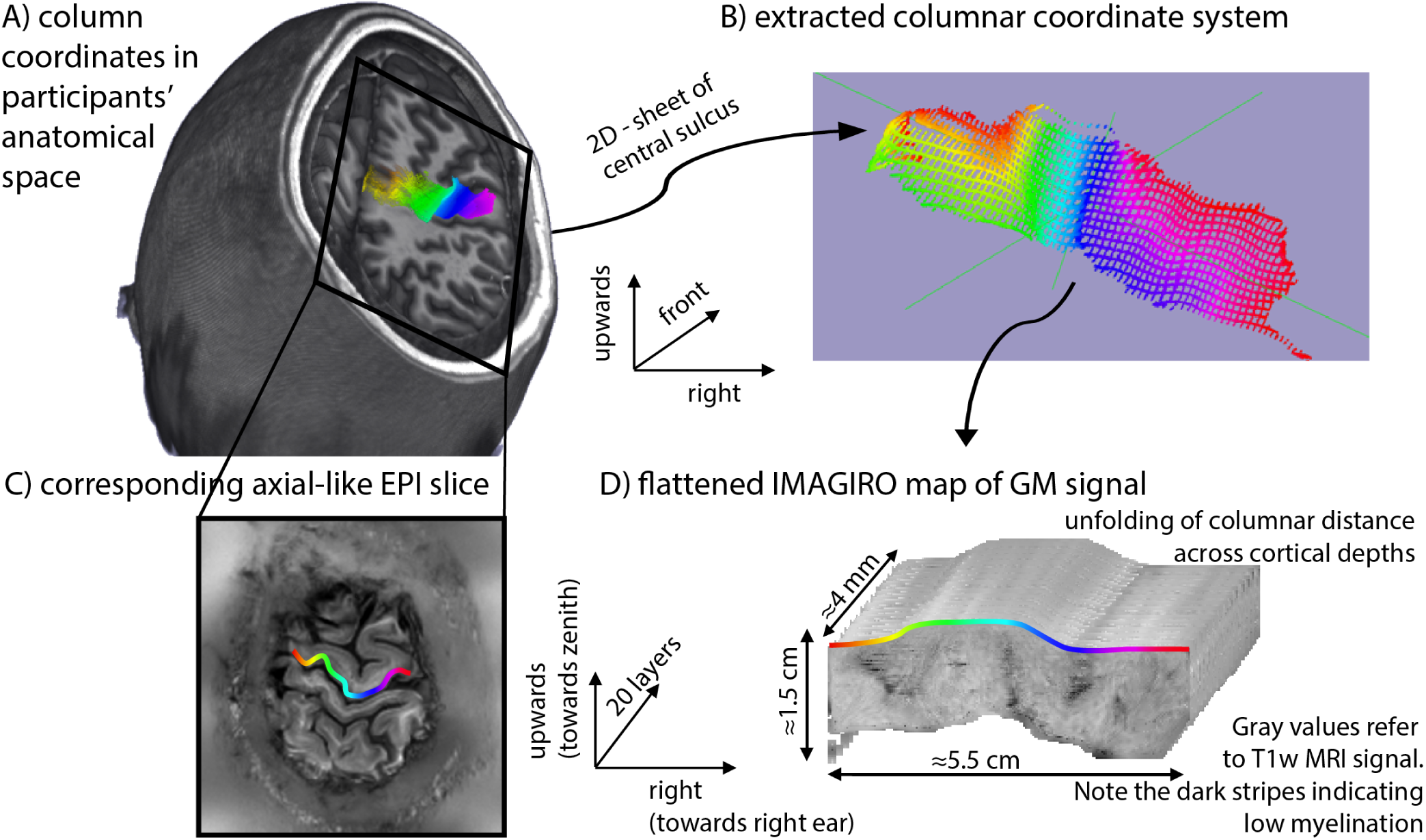
Schematic depiction of the unfolding method, IMAGIRO. IMAGIRO (origami written backwards) flattens the participants’ folded cortex (A, B) to a 3D-rectangular matrix (D). The colorful grid refers to a coordinate system of columnar distances across the depth of the central sulcus (vertical lines) and across the medial-lateral direction (horizontal lines). The lines are 1 mm apart, which corresponds to about five estimated column structures. The colors refer the relative medial-lateral position of the central sulcus (not to be confused with finger preference). In the IMAGIRO view, different body parts are represented from left to right. The depth of the central sulcus refers to the vertical direction. This flattened map can be obtained for different cortical depths (dimension into the page). (C) depicts the orientation of the EPI acquisition in this space for reference.

**Figure S4.**
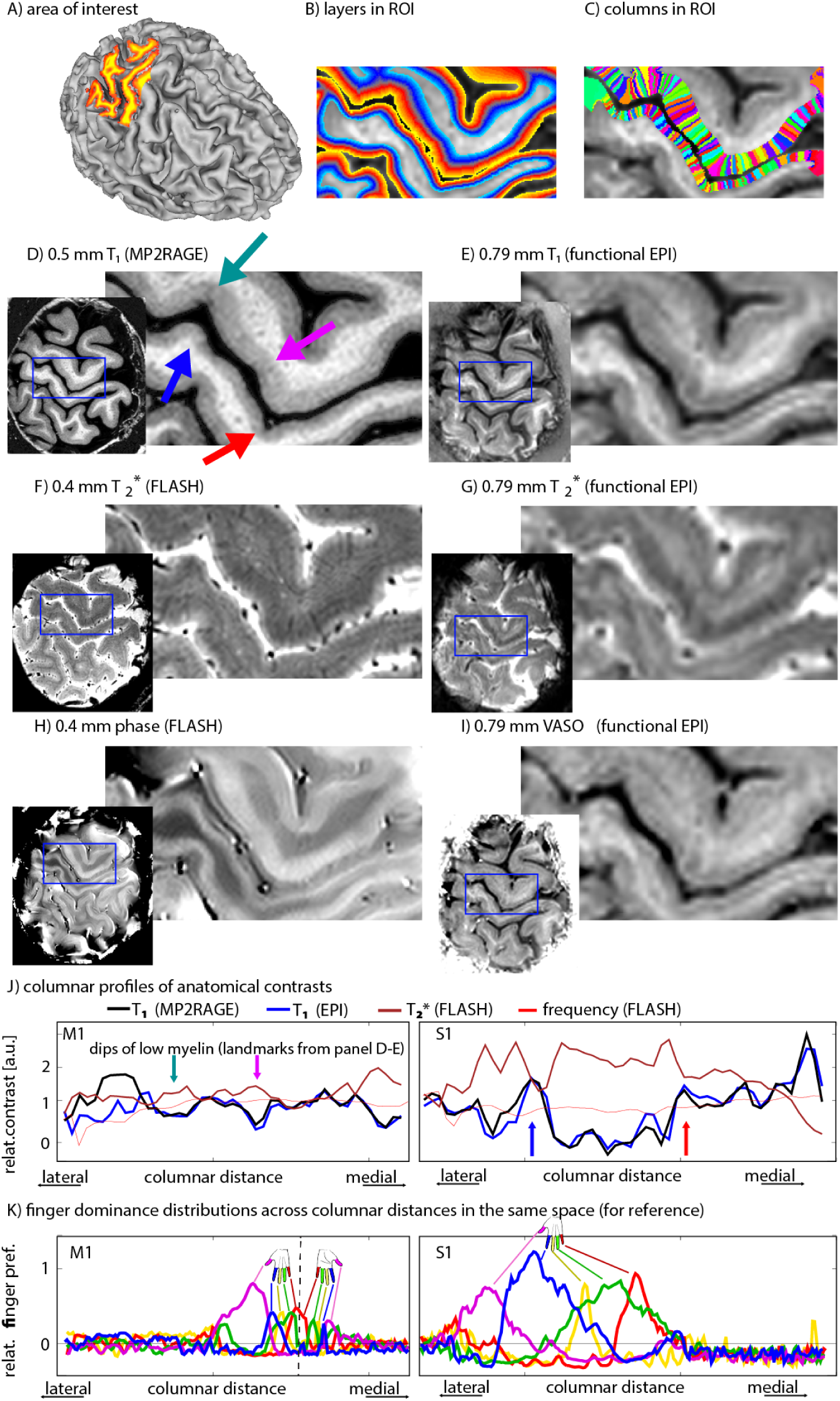
Anatomical contrasts used in this study. A) presents visual guidance to the following panels. The region of interest is highlighted in red. B-C) display the cortical coordinate system of layers and columns in the investigated area. Note that the size of layer and column structures are smaller than the effective resolution of 0.79 mm. They are estimated in an upscaled space. D) shows the 0.5 mm quantitative T1-MP2RAGE UNI contrasts. E) depicts T1-weighted contrast of the functional data. This contrast is used to identify GM/WM and GM/CSF boundaries for cortical depth estimations. F) demonstrates a 0.4 mm quantitative T2* map. This contrast is used to identify voxels of large draining veins. G) exhibits the BOLD EPI intensity. Similar structures and vessels can be seen as in panel F. H) shows a 0.4 mm quantitative FLASH phase map. I) depicts the average VASO intensity. The anatomical contrasts shown in the panels of the figure are used as reference and to find the position of body-part boundaries (panel K).

The repeatability of the double-hand representations and the their position across the depth of the central sulcus and compared to the action patches has also been investigated in test-retest experiments across days in the same participants with the same tasks. Fig. S6 depicts that the position of the double-hand representation is highly reproducible across days.

### Time courses

The grasping vs. retraction task does not directly isolate the action patches shown in Figs 1-3, S8. It is important to point out that both patches in the primary motor cortex show increased activity during both action tasks. The individual patches could be identified based on their differential activity strengths only (Fig. S7).

### Resting-state functional connectivity along the lateral position across the central sulcus

We used resting-state functional connectivity to investigate the organization principle across the lateral position of the central sulcus and the depth of the central sulcus. This is done to confirm the task results shown in Fig. 3 with independent resting-state data. Topographical patterns of functional connectivity were investigated by means of resting-state connectivity kernels. In this method, the correlation of the fluctuations of any given voxel is compared to its surroundings. Then, the spatial pattern of the resting-state correlation are averaged for every voxel in M1. This procedure is done across superficial and deeper layers (Fig. S8A) in the IMAGIRO coordinate system. We find that the correlation within columns is larger than within layers (purple arrows in Fig. S8B-C). This confirms that the topographical representation alignment is more columnar organized than laminar organized. We find that the overall connectivity is not circular across the columnar dimensions. Instead, columnar structures have a higher functional connectivity across the depth of the central sulcus than across the lateral dimension (Blue ellipses have larger height than width in Fig. S8B-C). This confirms that there is a higher connectivity within body part representations than across body part representations. We find that deeper layers show slight deviations from an exponential decay of functional connectivity across columnar distance. Instead, we find a reduction of functional connectivity in a center-surround pattern. This is despite the fact that deeper layers have larger lateral vessels than superficial layers (Fig. S8D).

The fact that deeper layers show less overlap of neighboring body parts confirms the task results shown in Fig. 3C. One potential origin (among others) of the sharper body part representation might be associated with previously described phenomena of surround inhibition in M1.

### Variability of results across the depth of the central sulcus

It is important to note that all of the results shown in Figs. 1-4, S1-S10 refer to the Brodmann area BA4a which is the superior portion of M1 on the precentral bank of the central sulcus close to the crown of the precentral gyrus. This part of the primary motor cortex is not to be confused with the Brodmann area BA4p, which is located deeper within the central sulcus and contains most of the cortio-motorneuron cells (Rathelot and Strick 2006, 2009; Lemon 2008). It has been shown with PET that both BA4p and BA4a represent movements of the fingers (Geyer 1996). It should be noted, that the double-hand representation found in this study does not refer to the two hand representations across BA4a and BA4p. Fig. S9 depicts the alignment of the respective sub areas of the primary motor cortex.

### Pilot experiments to determine the cortical hemisphere with the optimal sensitivity

Five participants were invited to participate in pilot experiments to identify whether it is optimal to focus on the left or the right hand and hemisphere. We found that due to the non-spherical shape of human head, the 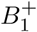-fields are higher in the right motor cortex compared to the left motor cortex (Fig. S10). This effect is due to the fact that at 300 MHz, the period of oscillation is comparable to the time that it takes for the radio-field to travel with the speed of light through the head, resulting in destructive interference. Without a parallel transmit system available, we thus decided to focus on the right motor cortex during left hand finger movements.

**Figure S5.**
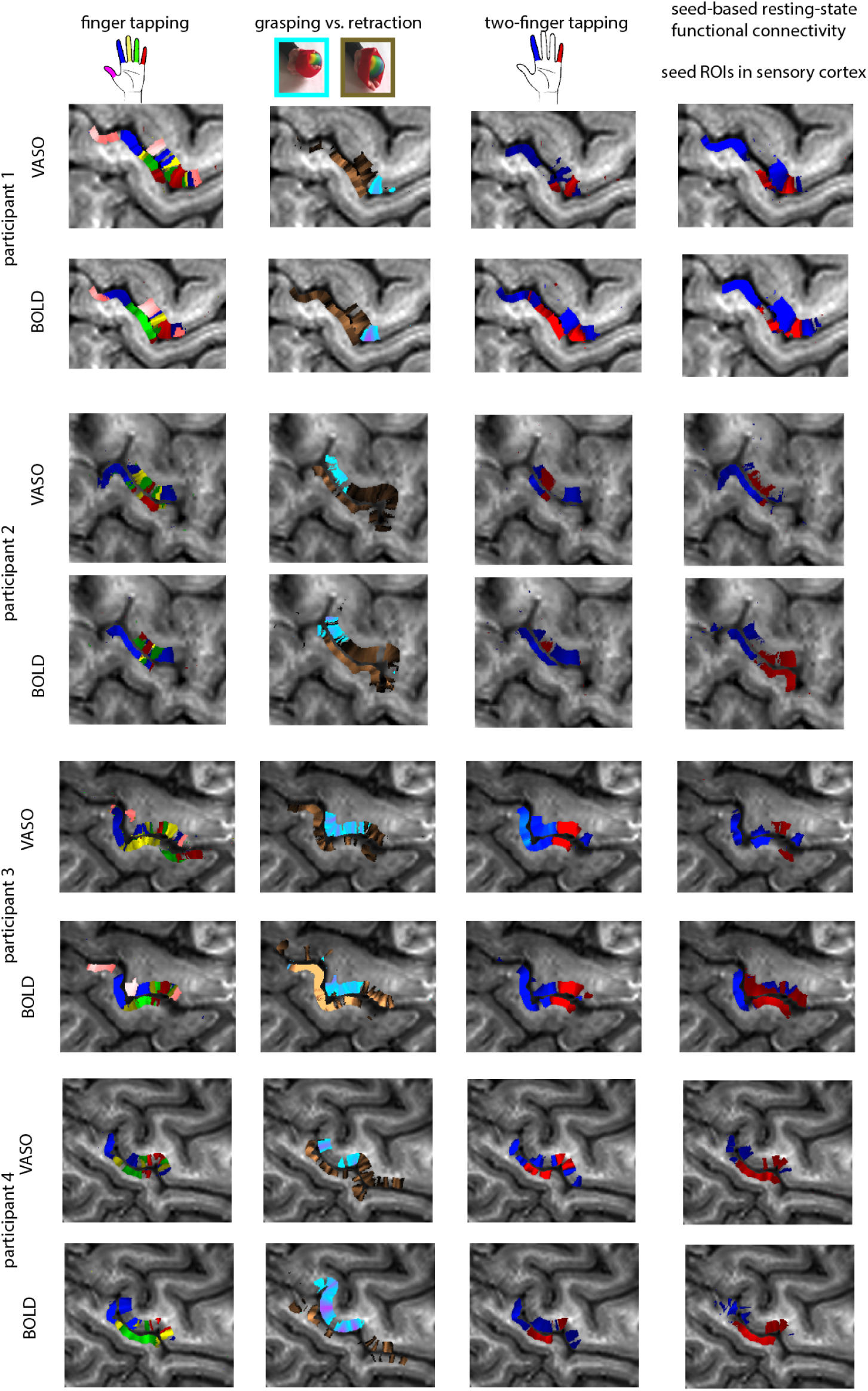
Robustness of the digit representations across contrasts and tasks. The maps depicted confirm the reproducibility of the maps shown in Fig. 2,4. Across all contrasts and tasks, multiple mirrored representations of individual fingers can be seen.

**Figure S6.**
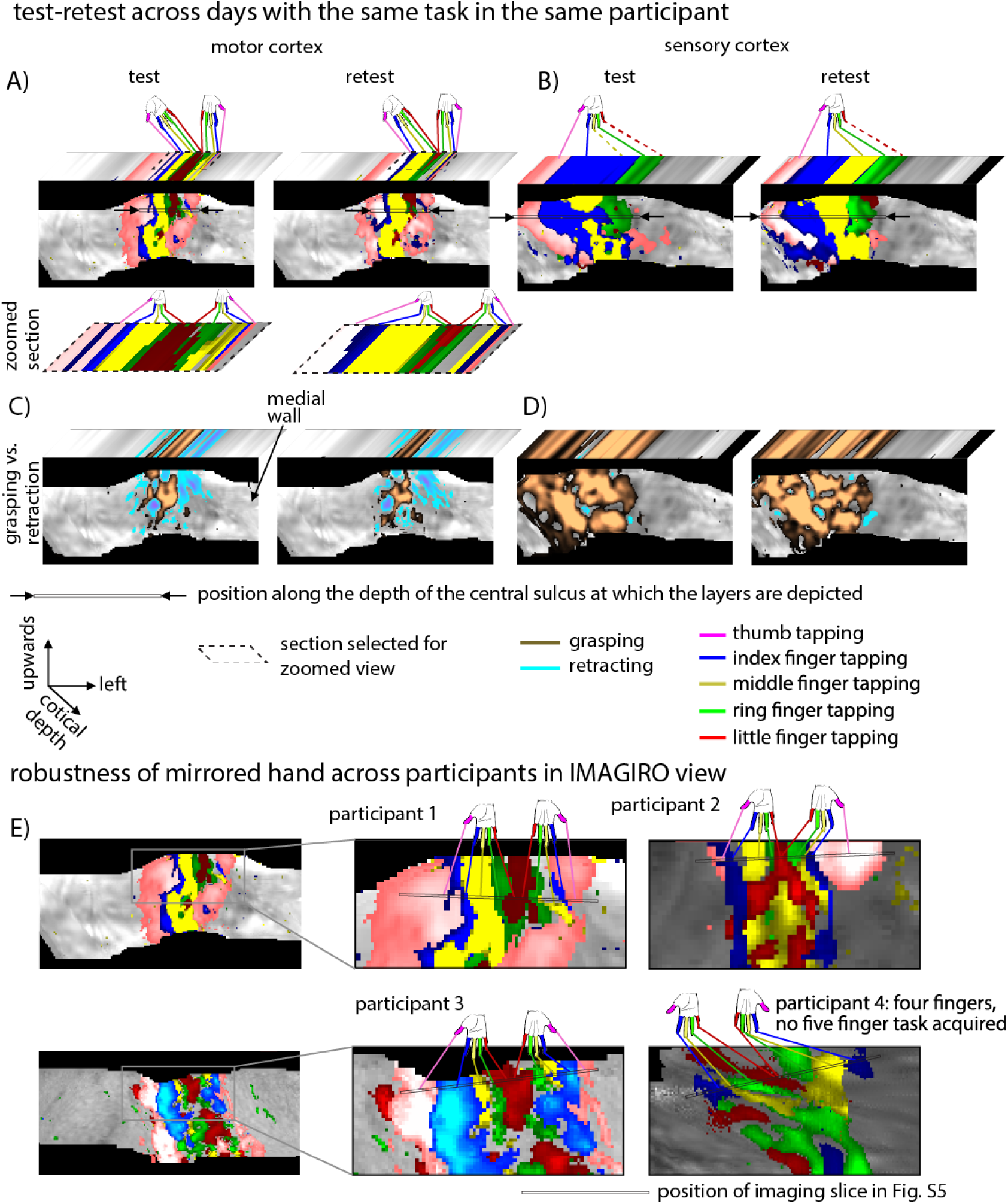
Test-retest results of the double-hand representation with the same participant and the same tasks across different days. Every activation map shown here refers to an independent experiment of 68 min (two times 34 min) on one day. The IMAGIRO maps depict the activation of the middle cortical layer across the entire central sulcus. At one position along the depth of the central sulcus (black arrows), the activation is shown across all cortical depths. A) It can be seen that the double-hand representation is reproducible across days. B) For the sake of comparison of the hand representation size, the results of the sensory cortex are shown as well. Panels C)-D) depict the corresponding grasping vs. retraction preferences. In the motor cortex it can be seen that the two hand representations refer to different action preferences. Panel E) depicts the mirrored finger representation across multiple participants. These are the same data of Fig. S5 depicted in the IMAGIRO view.

**Figure S7.**
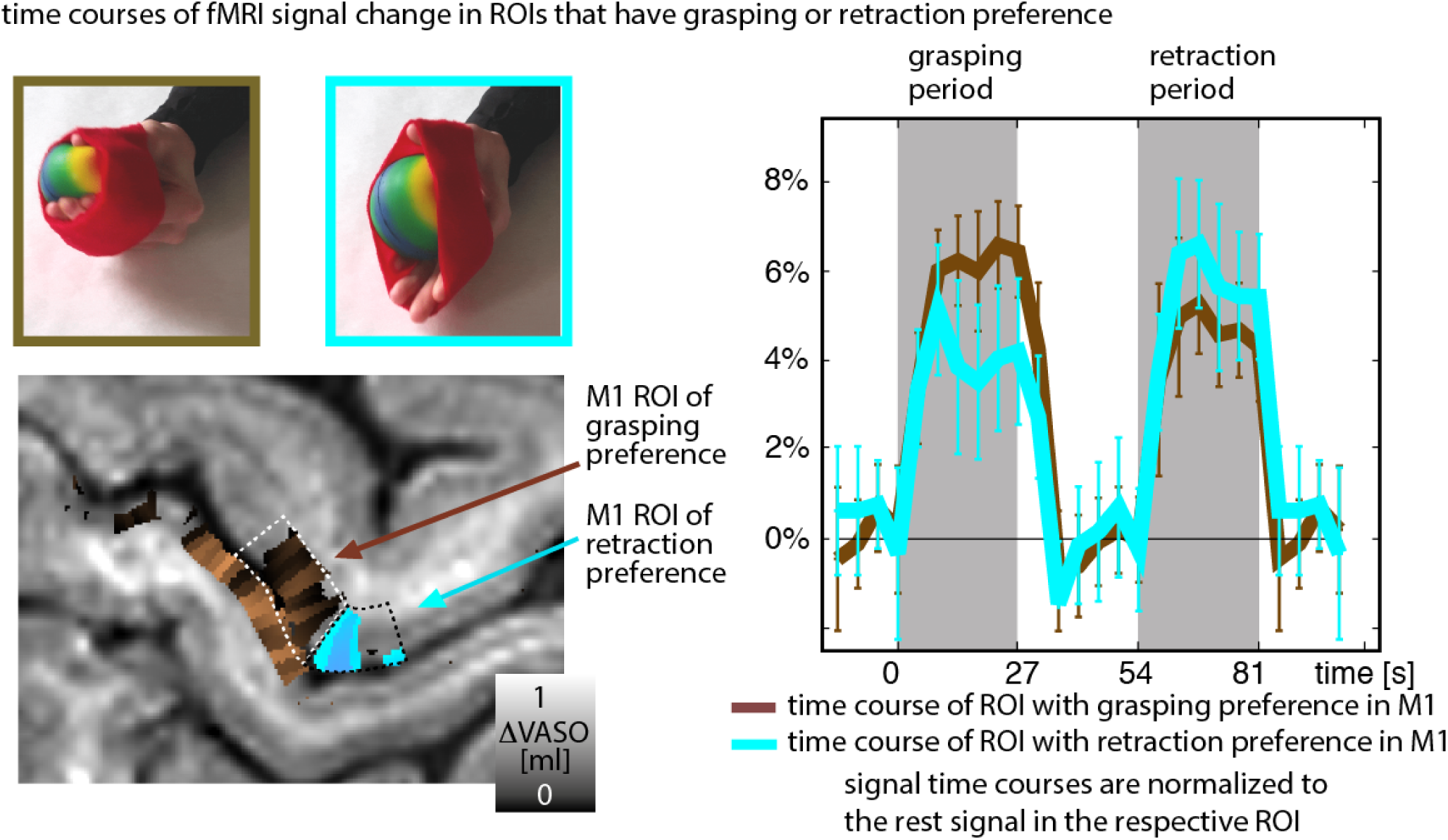
Time courses grasping vs. retraction task. Data refer to averages across all trials in two 34 min experiments. The error-bars refer to the standard deviation across trials. The time course shows that both action patches (retraction and the grasping patch) become active during both motor tasks. Their respective activity, however, is modulated based on the specific motor action conducted.

**Figure S8.**
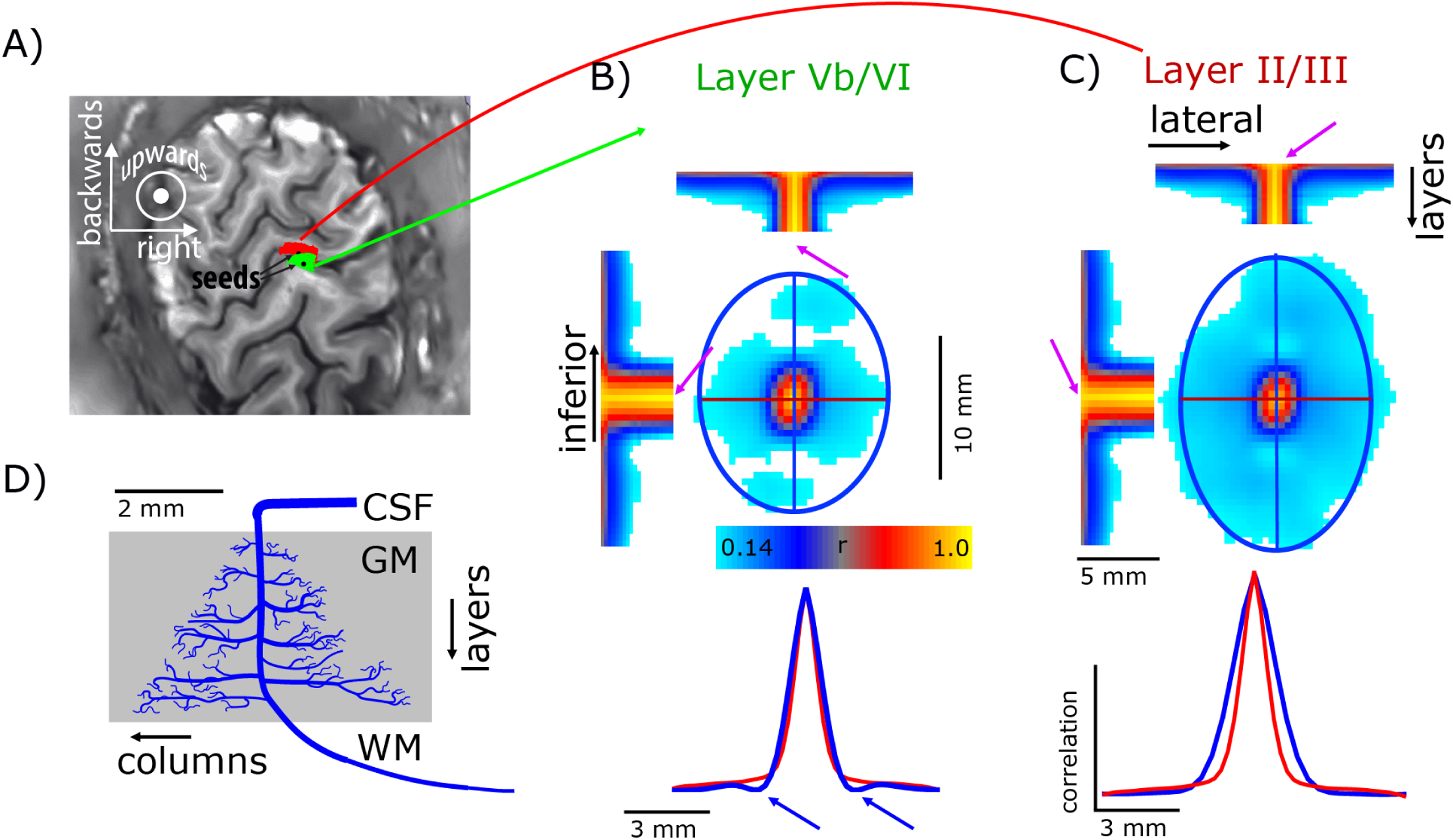
Topography of functional connectivity measured across cortical depth by means of connectivity kernels. Panel A) schematically depicts one iteration of a local connectivity estimation. For any seed (black dot) the functional connectivity is estimated for its surrounding (green and red areas respectively). The mean connectivity kernels are shown for deeper and superficial layers in panels B-C). The red and blue lines refer to cross-sections of the kernel along the central sulcus depths and along the lateral position. The result in this figure shows that there is a higher functional connectivity within body part representation than across body part representations (blue outline is a vertical ellipse rather than a circle). We find that deeper layers show less overlap of body parts than the upper layers (side lobes at blue arrows in panel B). This is visible despite the larger potential vascular bias in deeper layers compared to superficial layers, as indicated by the vein drawing shown in panel D (reproduced from Duvernoy 1981).

**Figure S9.**
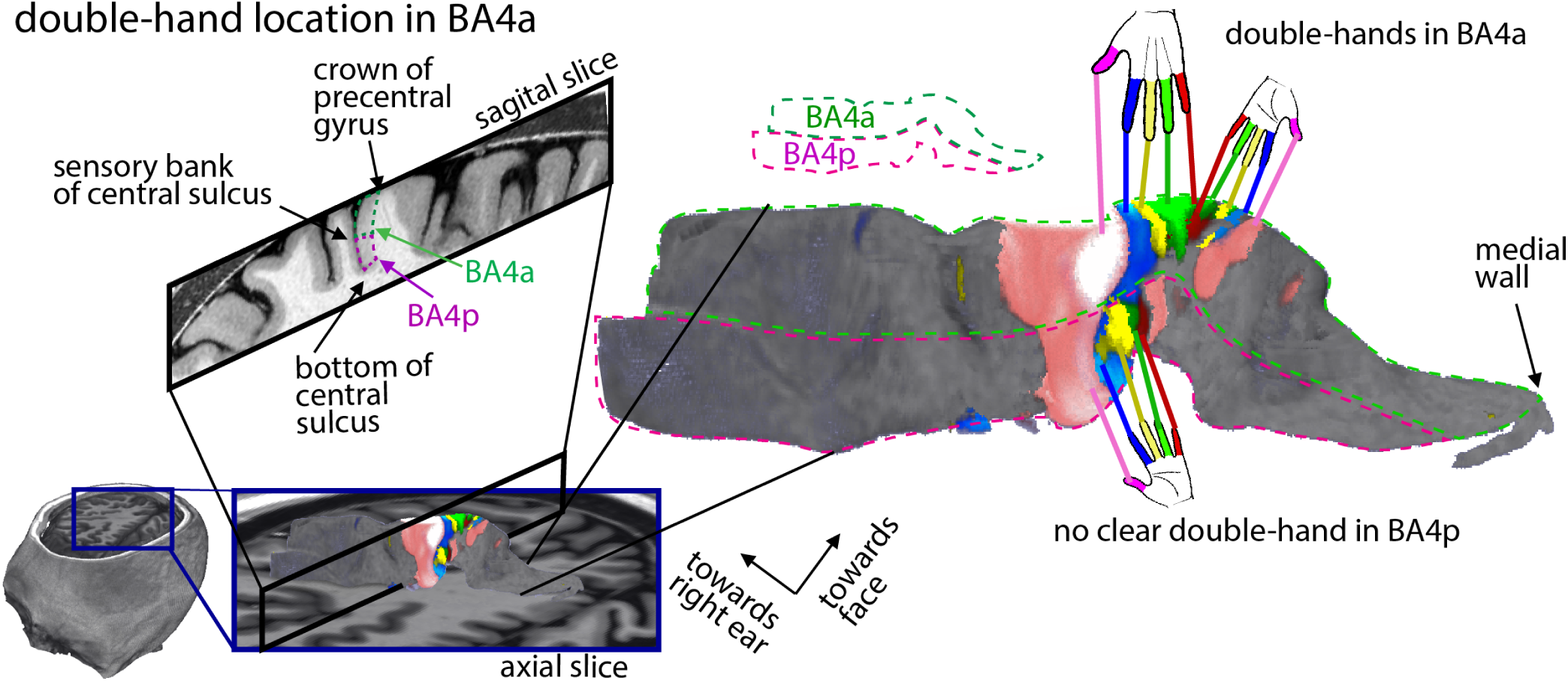
Position of the double-hand representation across the sub areas BA4p and BA4a. The double-hand representation in a mirrored finger pattern found in this study is located with in BA4a. Deeper in the central sulcus, in BA4p, an additional hand representation can be found. The hand representation in BA4p is roughly aligned with the superior hand representation in BA4a without a clear mirrored pattern.

**Figure S10.**
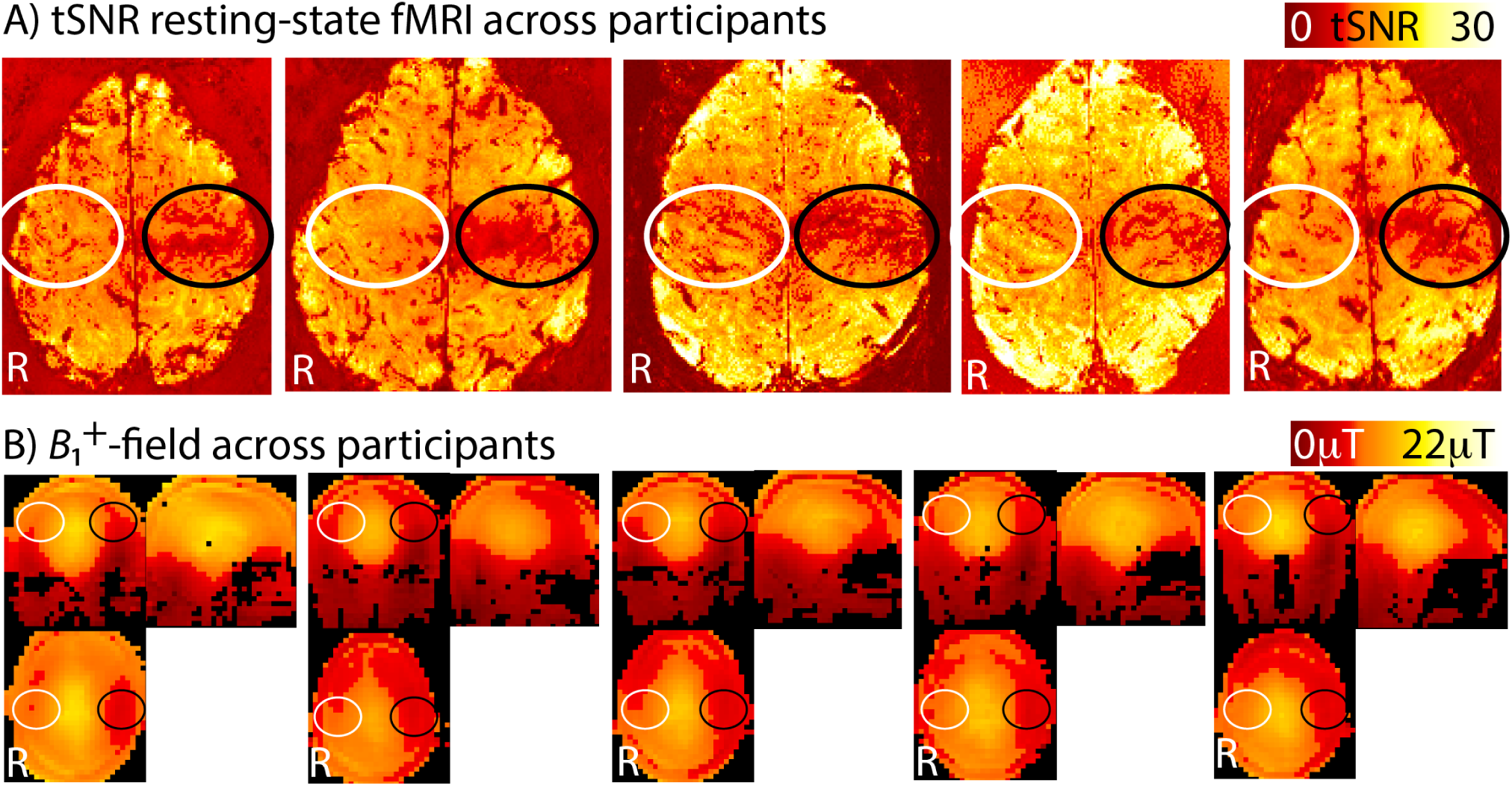
Comparison of sequence optimization of the left and the right sensorimotor cortex. A) tSNR maps of the functional VASO contrast show that the right sensorimotor cortex (white ellipses) has a clearly higher signal stability than the left sensorimotor cortex (black ellipses). This made us choose the right sensorimotor cortex during left-hand motor tasks in this study. B) We believe the lower sensitivity in the left sensorimotor cortex is due to asymmetric RF-interference artefacts in the presence of a non-spherical object (human head) in the RF-coil.

## Footnotes

1 Note, that in the context of fMRI, ‘columnar’ and ‘laminar’ do not refer to cytoarchitectonically and myelo-architectonically defined layers and receptive field defined columns (see discussion in the supplementary information 6). Instead they refer to mesoscopic structures that are aligned along (tangential) and perpendicular (radial) to the cortical depths. Also known as “cortical depths” and “hypercolumns”.

2 Such numbers of participants are considered reasonable even from the more conservative estimates (http://www.russpoldrack.org/2018/04/how-can-one-do-reproducible-science.html), as long as every participant is investigated for more than 4 hours (e.g., in visual neuroscience).

## Bibliography

Beck, S., and Hallett, M. (2011). Surround inhibition in the motor system. Exp. Brain Res. 210, 165–172.

Chainay, H., Krainik, A., Tanguy, M., Gerardin, E., and Bihan, D. Le (2004). Foot, face and hand representation in the human supplementary motor area. Neuroreport 15, 1–5.

Cheney, P.D., and Fetz, E.E. (1985). Comparable patterns of muscle facilitation evoked by individual corticomotoneuronal (CM) cells and by single intracortical microstimuli in primates: evidence for functional groups of CM cells. J. Neurophysiol. 53, 786–804.

d’Avella, A., and Bizzi, E. (2005). Shared and specific muscle synergies in natural motor behaviors. Proc. Natl. Acad. Sci. 102, 3076–3081.

Dechent, P., and Frahm, J. (2003). Functional somatotopy of finger representations in human primary motor cortex. Hum. Brain Mapp. 18, 272–283.

De Martino, F., Moerel, M., Ugurbil, K., Goebel, R., Yacoub, E., and Formisano, E. (2015). Frequency preference and attention effects across cortical depths in the human primary auditory cortex. Proc. Natl. Acad. Sci. 112, 16036–16041.

Ding, S.L., Royall, J.J., Sunkin, S.M., Ng, L., Facer, B.A.C., Lesnar, P., Guillozet-Bongaarts, A., McMurray, B., Szafer, A., Dolbeare, T.A., et al. (2016). Comprehensive cellular-resolution atlas of the adult human brain. J. Comp. Neurol. 524, 3127–3481.

Duvernoy, H.M., Delon, S., and Vannson, J.L. (1981). Cortical blood vessels of the human brain. Brain Res. 7, 519–579.

Ejaz, N., Hamada, M., and Diedrichsen, (2015). Hand use predicts the structure of representations in sensori-motor cortex. Nat. Neurosci. 103, 1–10.

Fischl, B., and Dale, A.M. (2000). Measuring the thickness of the human cerebral cortex from magnetic resonance images. Proc. Natl. Acad. Sci. 97, 11050–11055.

Fracasso, A., Petridou, N., and Dumoulin, S.O. (2016). Systematic variation of population receptive field properties across cortical depth in human visual cortex. Neuroimage 139, 427–438.

Geyer, S., Ledberg, A., Schleicher, A., Kinomura, S., Schormann, T., Burgel, U., Klingberg, T., Larsson, J., Zilles, K., and Roland, P.E. (1996). Two different areas within the primary motor cortex of man. Nature 382, 805–807.

Graziano, M.S.A., Taylor, C.S.R., and Moore, T. (2002). Complex movements evoked by micro-stimulation of precentral cortex. Neuron 34, 841–851.

Graziano, M.S.A. (2016). Ethological Action Maps: A Paradigm Shift for the Motor Cortex. Cel 20, 121–132.

Hatsopoulos, N.G. (2010). Columnar organization in the motor cortex. Cortex 46, 270–271.

Hlustík, P., Solodkin, A., Gullapalli, R.P., Noll, D.C., and Small, S.L. (2001). Somatotopy in human primary motor and somatosensory hand representation revisited. Cereb. Cortex 11, 312–321.

Hua, J., Jones, C.K., Qin, Q., and van Zijl, P.C.M. (2013). Implementation of vascular-space-occupancy MRI at 7T. Magn. Reson. Med. 69, 1003–1013.

Hubel, D.H., Wiesel, T.N., (1968). Receptive fields and functional architecture of monkey striate cortex. J. Physiol. 195, 215–243.

Hubel, D.H., Wiesel, T.N., 1972. Laminar and columnar distribution of geniculo-cortical fibers in the macaque monkey. J. Comp. Neurol. 146, 421–450.

Huber, L., Ivanov, D., Krieger, S.N.S.N., Streicher, M.N.M.N., Mildner, T., Poser, B.A.B.A., Möller, H.E., Turner, R., Moller, H.E., and Turner, R. (2014). Slab-selective, BOLD-corrected VASO at 7 tesla provides measures of cerebral blood volume reactivity with high signal-to-noise ratio. Magn. Reson. Med. 72, 137–148.

Huber, L., Goense, J., Kennerley, A. J., Trampel, R., Guidi, M., Reimer, E., Turner, R., Möller, H. E. (2015a). Cortical lamina-dependent blood volume changes in human brain at 7T. NeuroImage, 107, 23–33. http://doi.org/10.1016/j.neuroimage.2014.11.046

Huber, L., Goense, J. B. M., Kennerley, A. J., Guidi, M., Trampel, R., Turner, R., Möller, H. E. (2015b). Micro-and macrovascular contributions to layer-dependent blood blood volume fMRI: A multi-modal, multi-species comparison. In Proceedings of the International Society of Magnetic Resonance in Medicine (Vol. 23, p. 2114). http://doi.org/10.7490/f1000research.1114804.1

Huber, L. (2015c). Mapping human brain activity by functional magnetic resonance imaging of blood volume. Thesis of the University of Leipzig. http://nbn-resolving.de/urn:nbn:de:bsz:15-qucosa-165252

Huber, L., Handwerker, D.A., Jangraw, D.C., Chen, G., Hall, A., Stüber, C., Gonzalez-Castillo, J., Ivanov, D., Marrett, S., Guidi, M., et al. (2017). High-resolution CBV-fMRI allows mapping of laminar activity and connectivity of cortical input and output in human M1. Neuron 96, 1253–1263.

Huber, L., Tse, D.H.Y., Wiggins, C.J., Jangraw, D.C., Bandettini, P.A., Poser, B.A., and Ivanov, D. (2018). Ultra-high resolution blood volume fMRI and BOLD fMRI in humans at 9.4T: Capabilities and challenges. Neuroimage ahead of print.

Hurley, A.C., Al-Radaideh, A., Bai, L., Aickelin, U., Coxon, R., Glover, P., and Gowland, P.A. (2010). Tailored RF pulse for magnetization inversion at ultrahigh field. Magn. Reson. Med. 63, 51–58.

Indovina, I., and Sanes, J.N. (2001). On somatotopic representation centers for finger movements in human primary motor cortex and supplementary motor area. Neuroimage 13, 1027–1034.

Ivanov, D., Barth, M., Uludag, K., and Poser, B.A. (2015). Robust ACS acquisition for 3D echo planar imaging. In Proceedings of the International Society of Magnetic Resonance in Medicine, p. 2059.

Kennerley, A.J., Berwick, J., Martindale, J., Johnston, D., Papadakis, N.G., and Mayhew, J.E. (2005). Con-current fMRI and optical measures for the investigation of the hemodynamic response function. Magn. Reson. Med. 54, 354–565.

Kennerley, A. J., Huber, L., Mildner, T., Mayhew, J., Turner, R., Moeller, H. E., Berwick, J. (2013). Does VASO contrast really allow measurement of CBV at High Field (>=7T)? An in-vivo quantification using concurrent Optical Imaging Spectroscopy. In Proc Intl Soc Mag Reson Med (p. 0757).

Kim, S.-G., and Ogawa, S. (2012). Biophysical and physiological origins of blood oxygenation level-dependent fMRI signals. J. Cereb. Blood Flow Metab. 32, 1188–1206.

Klein, B.P., Fracasso, A., van Dijk, J.A., Paffen, C.L.E., te Pas, S.F., and Dumoulin, S.O. (2018). Cortical depth dependent population receptive field attraction by spatial attention in human V1. Neuroimage 176, 301–312.

Kolasinski, J., Makin, T.R., Jbabdi, S., Clare, S., Stagg, C.J., and Johansen-Berg, H. (2016). Investigating the Stability of Fine-Grain Digit Somatotopy in Individual Human Participants. J. Neurosci. 36, 1113–1127.

Kuehn, E., Dinse, J., Jakobsen, E., Long, X., Schäfer, A., Bazin, P.-L., Villringer, A., Sereno, M.I., and Mar- gulies, D.S. (2017). Body Topography Parcellates Human Sensory and Motor Cortex. Cereb. Cortex 27, 3790–3805.

Kwan, H.C., MacKay, W.A., Murphy, J.T., and Wong, Y.C. (1978). Spatial organization of precentral cortex in awake primates. II. Motor outputs. J. Neurophysiol. 41, 1120–1131.

Lemon, R.N. (2008). Descending Pathways in Motor Control. Annu. Rev. Neurosci. 31, 195–218.

Lu, H., Golay, X., Pekar, J.J., and van Zijl, P.C.M. (2003). Functional magnetic resonance imaging based on changes in vascular space occupancy. Magn. Reson. Med. 50, 263–274.

Makin, T. R., Ejaz, N., Wesselink, D. B., Tarall-Jozwiak, A., van den Heiligenberg, F. M., Cardinali, L., … Diedrichsen, J. (2019). Obtaining and maintaining cortical hand representation as evidenced from acquired and congenital handlessness. ELife, 8, 1–19. http://doi.org/10.7554/elife.37227

Marques, J.P., Kober, T., Krueger, G., van der Zwaag, W., Van de Moortele, P.-F., and Gruetter, R. (2010). MP2RAGE, a self bias-field corrected sequence for improved segmentation and T1-mapping at high field. Neuroimage 49, 1271–1281.

Meier JD, Aflalo TN, Kastner S, Graziano MS. Complex organization of human primary motor cortex: a high-resolution fMRI study. J Neurophysiol. 2008 Oct;100(4):1800–12. doi:10.1152/jn.90531.2008. Epub 2008 Aug 6

Menon, R.S. (2002). Postacquisition suppression of large-vessel BOLD signals in high-resolution fMRI. Magn. Reson. Med. 47, 1–9.

Omrani, M., Kaufman, M.T., Hatsopoulos, N.G., and Cheney, P.D. (2017). Perspectives on classical controversies about the motor cortex. J. Neurophysiol. jn.00795.2016.

Olman, C., Pickett, K. a, Schallmo, M.P., and Kimberley, T.J. (2012). Selective BOLD responses to individual finger movement measured with fMRI at 3T. Hum. Brain Mapp. 33, 1594–1606.

Park, M.C., Belhaj-Saïf, A., Gordon, M., and Cheney, P.D. (2001). Consistent features in the forelimb representation of primary motor cortex in rhesus macaques. J. Neurosci. 21, 2784–2792.

Penfield, W., and Boldrey, E. (1937). Somatic Motor and Sensory Representation in Man. Brain 389–443.

Porter, R., and Lemon, R. (2012). Corticospinal function and voluntary movement (Oxford University Press).

Poser, B.A., Koopmans, P.J., Witzel, T., Wald, L.L., and Barth, M. (2010). Three dimensional echo-planar imaging at 7 tesla. Neuroimage 51, 261–266.

Rathelot, J.-A., and Strick, P.L. (2006). Muscle representation in the macaque motor cortex: An anatomical perspective. Proc. Natl. Acad. Sci. 103, 8257–8262.

Rathelot, J.-A., and Strick, P.L. (2009). Subdivisions of primary motor cortex based on corticomotoneuronal cells. Proc. Natl. Acad. Sci. 106, 918–923.

Sanes, J.N., Donoghue, J.P., Thangaraj, V., Edelman, R.R., and Warach, S. (1995). Shared neural substrates controlling hand movements in human motor cortex. Science 268, 1775–1777.

Schellekens, W., Petridou, N., and Ramsey, N.F. (2018). Detailed somatotopy in primary motor and somatosensory cortex revealed by Gaussian population receptive fields. Neuroimage, Press. 179, 337–347.

Schellekens, W., Petridou, N., and Ramsey, N.F. (2017). Mapping the human homunculus with receptive field analysis. In Society for Neuroscience, p. 62.13.

Schieber, M.H., and Hibbard, L.S. (1993). How somatotopic is the motor cortex hand area? Science (80). 261, 489–492.

Schieber, M.H. (2002). Motor cortex and the distributed anatomy of finger movements. Adv. Exp. Med. Biol. 508, 411–416.

Schluppeck, D., Sanchez-Panchuelo, R.M., and Francis, S.T. (2017). Exploring structure and function of sensory cortex with 7 T MRI. Neuroimage ahed of print.

Siero, J.C.W., Hermes, D., Hoogduin, H., Luijten, P.R., Ramsey, N.F., and Petridou, N. (2014). BOLD matches neuronal activity at the mm scale: A combined 7T fMRI and ECoG study in human sensorimotor cortex. Neuroimage 101, 177–184.

Strick, P.L., and Preston, J.B. (1982). Two representations of the hand in area 4 of a primate. I. Motor output organization. J.Neurophysiol. 48, 139–149.

Strother, L., Medendorp, W.P., Coros, A.M., and Vilis, T. (2012). Double representation of the wrist and elbow in human motor cortex. Eur. J. Neurosci. 36, 3291–3298.

Sanchez Panchuelo, R.M., Ackerley, R., Glover, P.M., Bowtell, R.W., Wessberg, J., Francis, S.T., and Mc-Glone, F. (2016). Mapping quantal touch using 7 tesla functional magnetic resonance imaging and single-unit intraneural microstimulation. Elife 5, e12812.

Thakur, P.H., Bastian, A.J., and Hsiao, S.S. (2008). Multidigit Movement Synergies of the Human Hand in an Unconstrained Haptic Exploration Task. J. Neurosci. 28, 1271–1281.

Turner, R. (2002). How much cortex can a vein drain? Downstream dilution of activation-related cerebral blood oxygenation changes. Neuroimage 16, 1062–1067.

Wagstyl, K., Lepage, C., Bludau, S., Zilles, K., Fletcher, P.C., Amunts, K., and Evans, A.C. (2018). Mapping Cortical Laminar Structure in the 3D Big-Brain. Cereb. Cortex 1–12

Woolsey, T.A., Rovainen, C.M., Cox, S.B., Henegar, M.H., Liang, G.E., Liu, D., Moskalenko, Y.E., Sui, J., and Wei, L. (1996). Neuronal units linked to microvascular modules in cerebral cortex: Response elements for imaging the brain. Cereb. Cortex 6, 647–660.

